# The pentameric chloride channel BEST1 is activated by extracellular GABA

**DOI:** 10.1101/2024.11.22.624909

**Authors:** Swati Pant, Stephanie W. Tam, Stephen B. Long

**Affiliations:** Structural Biology Program, Memorial Sloan Kettering Cancer Center, 1275 York Avenue, New York, NY 10065, USA; Graduate Program in Biochemistry and Structural Biology, Cell and Developmental Biology, and Molecular Biology, Weill Cornell Medicine Graduate School of Medical Sciences, New York, USA; Graduate Program in Physiology, Biophysics, and Systems Biology, Weill Cornell Medicine Graduate School of Medical Sciences, New York, USA

**Keywords:** Chloride channel, GABA, bestrophin, activation, receptor

## Abstract

Bestrophin 1 (BEST1) is chloride channel expressed in the eye, central nervous system (CNS), and other tissues in the body. A link between BEST1 and the principal inhibitory neurotransmitter γ-aminobutyric acid (GABA) has been proposed. The most appreciated receptors for extracellular GABA are the GABA_B_ G-protein coupled receptors and the pentameric GABA_A_ chloride channels, both of which have fundamental roles in the CNS. Here, we demonstrate that BEST1 is directly activated by GABA. Through functional studies and atomic-resolution structures of human and chicken BEST1, we identify a GABA binding site on the channel’s extracellular side and determine the mechanism by which GABA binding induces opening of the channel’s central gate. This same gate is activated by intracellular [Ca^2+^], indicating that BEST1 is controlled by ligands from both sides of the membrane. The studies demonstrate that BEST1, which shares no structural homology with GABA_A_, is a GABA-activated chloride channel. The physiological implications of this finding remain to be studied.

**Significance Statement:** *γ*-aminobutyric acid (GABA) is the principal inhibitory neurotransmitter in the central nervous system. Extracellular GABA is primarily sensed by GABA_B_ G-protein coupled receptors, and GABA_A_ pentameric chloride channels. We show that the chloride channel bestrophin-1 (BEST1) is also activated by extracellular GABA. The application of GABA, but not glycine or other endogenous molecules we tested, markedly potentiates chloride currents through the channel. Structural studies combined with electrophysiology and other functional studies reveal the mechanism of GABA activation. GABA binding within the outer entryway of the channel allosterically controls its gate, ‘the neck’. Molecular interactions with GABA resemble those in GABA_A_ receptors, despite a lack of homology between the channels. The physiological significance of GABA-activation in BEST1 remains to be studied.

## Introduction

*γ*-aminobutyric acid (GABA) is the primary inhibitory neurotransmitter in the central nervous system (CNS) (1). GABAergic signaling is mediated by the GABA_A_ and the GABA_B_ families of receptors. GABA_B_ receptors are G-protein coupled receptors that signal to adenyl cyclases and potassium (K^+^) channels when activated by GABA binding (2). GABA_A_ receptors, on the other hand, are pentameric chloride (Cl^-^) channels that are directly stimulated by the binding of GABA to sites within the channel’s large extracellular domain (3–5). Nineteen types of GABA_A_ subunits assemble as heteropentamers to form a myriad of receptor subtypes with finely tuned functional properties and a range of GABA affinities. GABA binding stimulates opening of GABA_A_ channels through allosteric coupling to the transmembrane region that contains the gate of the ion pore (3). Opening of GABA_A_ channels generally results in Cl^-^ influx in mature neurons, leading to hyperpolarization that decreases the chance of action potential firing. GABA_A_ receptors respond to GABA over a broad range of concentrations; synaptic receptors are tuned to the millimolar levels of GABA found in synapses, while receptors present in extrasynpatic areas are sensitive to GABA in the micromolar range (6).

While they have no apparent sequence or structural similarity to GABA_A_ receptors, the bestrophin family of proteins (BEST1-4 in humans) assemble as pentamers and function as Cl^-^ channels (7–9). Bestrophin channels are activated by the binding of calcium (Ca^2+^) to their cytosolic side (7, 8, 10). Mutations in human BEST1 cause degenerative eye diseases, which are typically characterized by structural changes in the macula, loss of central vision, and reduction of the light peak of the electrooculogram, which indicates abnormal RPE function (11–13). Among the mutations in BEST1 that have been evaluated, most disrupt the electrophysiological properties of the channel, suggesting that perturbations in channel function are causative for eye disease (8, 14–17). In addition to the eye, there is evidence that bestrophins are expressed in the colon, pancreatic duct cells, airways, and in the CNS (18–24). In the CNS, for which there has been some debate about the extent of BEST1 expression (23, 24), it has been proposed that BEST1 may mediate low-level release of glutamate and GABA in astrocytes by the permeation of these molecules through the pore of the channel (22, 25). However, the breadth of physiological functions of bestrophin channels is largely unknown.

Bestrophin proteins are highly conserved among metazoan organisms. One of the most studied orthologs, chicken BEST1, shares 74% amino acid sequence identity to human BEST1 within the structured region of the channel (9). X-ray crystallographic studies of chicken BEST1, and subsequent structures of human BEST1 and BEST2, reveal that the channel is a pentamer, composed of five bestrophin subunits, and that it contains a single ion conduction pore along the symmetry axis (9, 26, 27). The pore has two constrictions: the ‘aperture’ at its cytosolic end, and the ‘neck’ within the transmembrane region (9). Electrophysiological and cryo-EM studies have shown that the neck functions as the Ca^2+^ controlled gate of the channel (10, 28). In the absence of Ca^2+^, the neck is closed. In the Ca^2+^-activation process, Ca^2+^ binding to a cytosolic site, the ‘Ca^2+^ clasp’, induces opening of the neck that permits the flow of ions (9, 10, 28).

The aperture comprises a short, narrow constriction of the pore that is lined by the sidechains at amino acid 205 (Val205 in chicken BEST1, Ile205 in human BEST1) from the five subunits of the channel. The aperture functions as a molecular sieve that limits the size of molecules that can pass through the channel (10, 28). The aperture adopts the same conformation in all structures of BEST1 determined to date and does not respond to changes in intracellular Ca^2+^ concentrations, suggesting that it does not function as a gate (9, 10, 26–29). Rather, the aperture governs permeability properties of the channel, making small anions, such as Cl^-^, more permeable and larger anions, such as glutamate, lowly permeable (10, 28, 30, 31).

In addition to being activated by cytosolic Ca^2+^ (with an EC_50_ ∼ 140 nM for human BEST1), the channel also undergoes inactivation whereby currents through the channel decrease over time (32, 33). The inactivation process is itself controlled by the intracellular Ca^2+^ concentration (32, 33). In human BEST1, micromolar levels of Ca^2+^ cause more rapid inactivation (time constant of ∼ 2 min), whereas slower inactivation is observed at lower (∼140 nM) Ca^2+^ concentrations (32). Studies have demonstrated that inactivation occurs due to the binding of C-terminally located peptide of BEST1 to a site on the cytosolic surface of the channel (27, 28, 32, 33). Binding of this ‘inactivation peptide’ induces neck closure of the channel. Unlike most other channels that undergo inactivation due to different molecular perturbations of the ion conduction pathway, one structural element – the neck – serves as both an activation and an inactivation gate.

We have previously determined structures of chicken BEST1 that represent the primary gating steps for both Ca^2+^-dependent activation and Ca^2+^-dependent inactivation (9, 28). These structures revealed that the neck can adopt two structural conformations, a closed conformation that prevents ion flow, and an open conformation that permits it. The channel thereby toggles between conductive and non-conductive conformations that are energetically biased by the activation and inactivation processes. The conformational landscape observed in the previous structures suggest that the closed conformation is energetically favored. In structural studies where the construct of chicken BEST1 used (spanning amino acids 1-405) contains the inactivation peptide, only the closed conformation of the neck was observed, even though Ca^2+^ was present at the Ca^2+^ clasp (9, 28). We were able to obtain the open conformation of the neck by removing the inactivation peptide (using a construct spanning amino acids 1-345) (28). Even in this case, only approximately 14% of the particles in the cryo-EM analysis represented the open conformation, and the remainder represented a Ca^2+^-bound closed conformation of the channel, suggesting that the closed conformation is energetically favorable.

Previous cryo-EM structures of human BEST1 revealed a similar pattern (27). When the inactivation peptide was included in the construct for cryo-EM analysis, two subtly different conformations of the neck were observed – a closed conformation analogous to that observed previously for chicken BEST1, and what was referred to as a ‘partially open’ conformation in which the neck-lining residues are slightly dilated from their closed conformation. Analogous to our studies of chicken BEST1, a fully open conformation was not observed for human BEST1 when the construct used for structural analysis included the inactivation peptide. As for chicken BEST1, an open conformation of the human channel was obtained by removing the inactivation peptide, and in this case resulted in essentially all the particles adopting an open conformation. The observation that the open conformation of BEST1 has only been observed by removal of the inactivation peptide raises a possibility that the neck may not be able to open fully when the inactivation peptide is present.

Here, we identify a link between GABA and BEST1. We find that the ion channel activity of BEST1 is activated by extracellular GABA. Through functional approaches and atomic-level cryo-EM structures, we determine that GABA binds to an allosteric site on the extracellular side of the channel that controls its gating. In addition to its roles in Ca^2+^-activation and inactivation, the neck also functions as the GABA-dependent gate.

## Results

### Purified BEST1 is activated by GABA

Data from other laboratories have suggested that the inhibitory neurotransmitter GABA may permeate through BEST1, perhaps in ionic form (25). We first evaluated a possible effect of GABA on BEST1 using a fluorescence-based flux assay, where flow of anions is detected by a decrease in fluorescence (Fig. 1A). As we previously reported for chicken BEST1 (9, 10), Cl^-^ readily permeates through the channel as detected by a time-dependent decrease in fluorescence (Fig. 1*B*). When the Cl^-^ in the assay (as 65 mM NaCl) is replaced by 65 mM GABA, no fluorescence decrease is observed. This indicates that GABA does not appreciably flow through the channel in anionic form. However, when both GABA and Cl^-^ are present, a dramatic decrease in fluorescence is observed (Fig. 1*B*). These results suggested that GABA may potentiate the flow of Cl^-^ through chicken BEST1. A titration of GABA, in the context of a fixed concentration of Cl^-^, reveals a dose response activation by GABA in the millimolar range (Fig. 1*C*).

**Fig 1.**
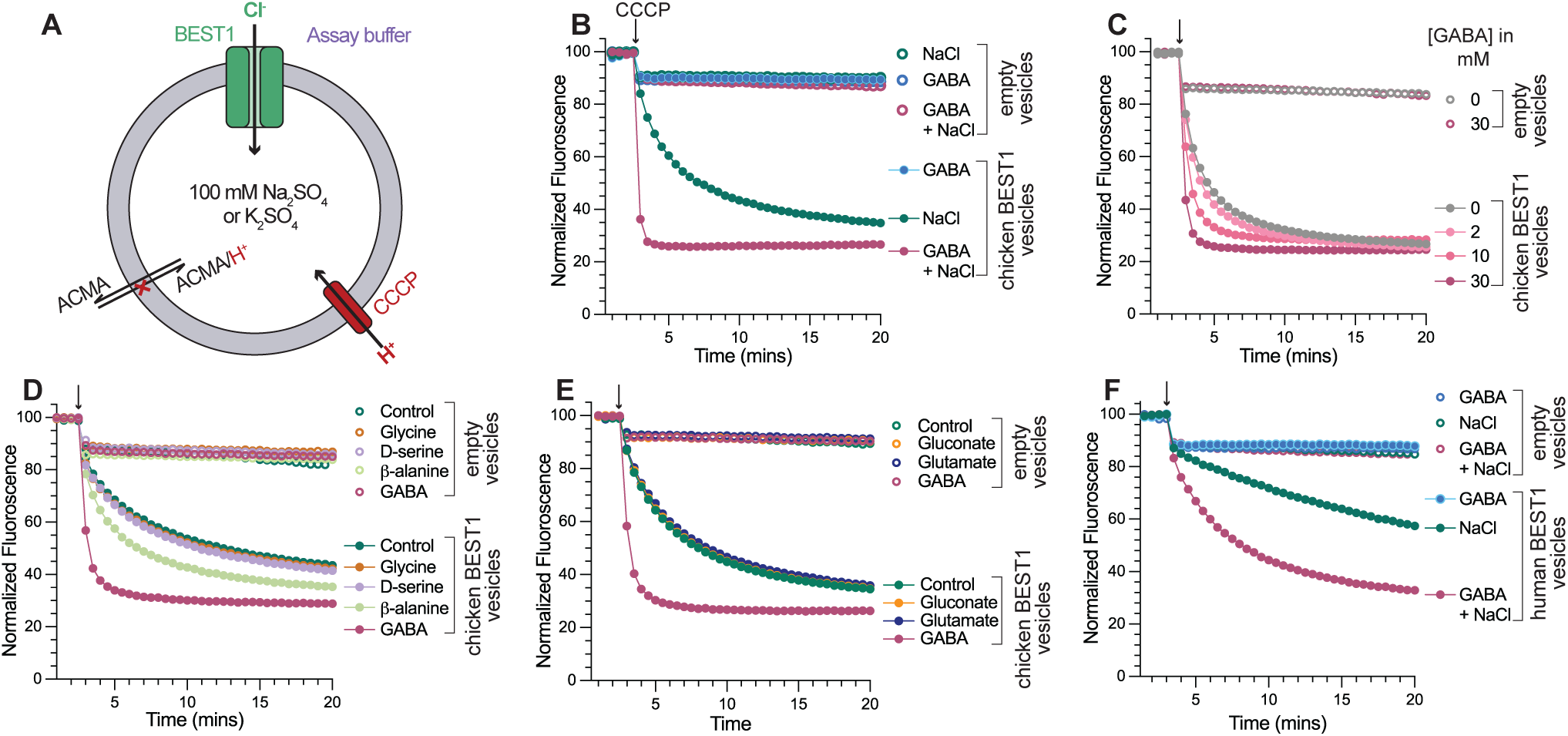
GABA augments Cl^-^ flow through BEST1. *(A)* Schematic of the flux assay. Purified BEST1 is reconstituted into liposomes. Liposomes either contain sodium sulphate (chicken BEST1) or potassium sulphate (human BEST1), which are not permeable through the channels. Liposomes are diluted in assay buffer containing NaCl and/or test compounds (Table S1). The flow of anions (Cl^-^) into the liposomes generates a negative potential, which is used to drive the uptake of protons through the ionophore CCCP. The protons quench the fluorescence of the pH sensitive dye ACMA. Thus, decrease in fluorescence is an indicator of anion flux through the channel. *(B)* Experiment testing the effect of GABA. The addition of GABA enhances Cl^-^ flux through chicken BEST1. On its own, GABA does not cause a decrease in florescence. Empty vesicles (i.e. vesicles without BEST1) are shown as controls. *(C)* GABA increases Cl^-^ flux through chicken BEST1 in a dose dependent manner. *(D-E)* Among endogenous molecules tested, only GABA dramatically enhances Cl^-^ flux. *(F)* Analogous experiment to *(B)* for human BEST1, showing that GABA potentiates Cl^-^ flux.

To evaluate how specific the response was to GABA, we used the flux assay to evaluate endogenous molecules with similar chemical structures. 30 mM concentrations of glycine, glutamate, gluconate, and D-serine had no discernable effect on the rate of fluorescence decrease (Fig. 1*D* and *E*). A slight potentiation of flux was observed for β-alanine, which has a highly similar chemical structure to GABA (Fig. 1*D* and Fig S1).

To evaluate if GABA has an analogous effect on the human channel, we purified human BEST1 and reconstituted it into liposomes for flux assay analysis. As observed using whole-cell electrophysiology (7, 18), but not previously in a reconstituted system, the assay indicated that Cl^-^ flows through purified human BEST1 (Fig 1*F*). We found that GABA accentuated the fluorescence decrease for the human BEST1, suggesting that GABA also potentiates Cl^-^ flow through the human channel (Fig. 1*F*). To further interrogate the effect of GABA, we studied human and chicken BEST1 using electrophysiology.

### Extracellular application of GABA activates human BEST1

We evaluated the effects of GABA on human BEST1 using whole-cell patch-clamp electrophysiology, with conditions based on previous electrophysiological studies of the channel (32). To minimize inactivation of BEST1, a low (10 nM) concentration of Ca^2+^ was used in the patch pipette. Mammalian cells (HEK293T) transfected with human BEST1 displayed characteristic BEST1 currents, with near-linear current-voltage (I-V) relationships (Fig. 2*D*). To evaluate a possible effect of GABA, repeated applications of 30 mM GABA were made by perfusing the extracellular solution followed by washout of the compound. The GABA applications resulted in corresponding periodic increases in ionic current (Fig. 2*C*). Analogous experiments using cells transfected only with green fluorescent protein (GFP), showed no appreciable ionic current or effect of GABA application (Fig. 2*B*).

**Fig 2.**
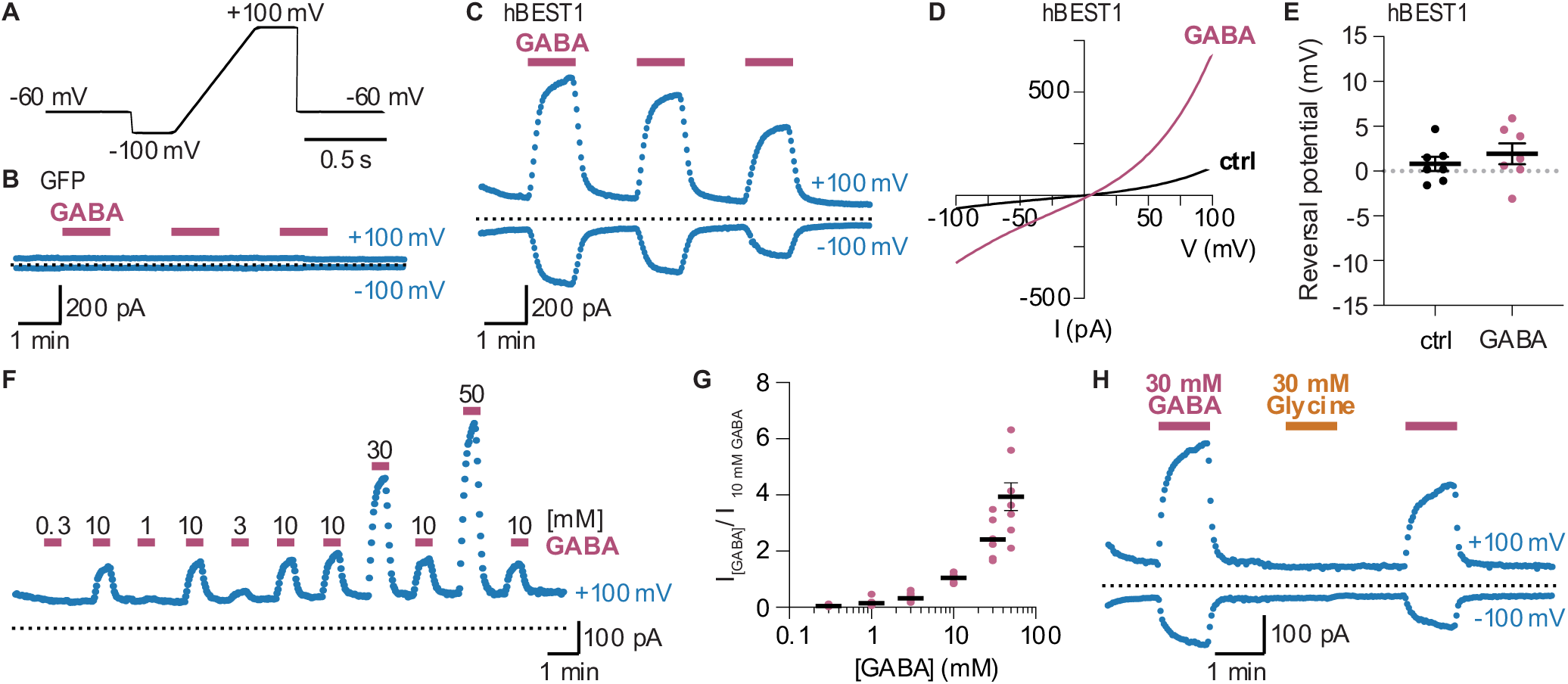
Whole-cell patch-clamp electrophysiology indicates that extracellular GABA activates human BEST1. *(A)* Voltage protocol using a train of 0.5 s voltage ramps from −100 mV to +100 mV (from a holding potential of −60 mV), repeated every 2 s. *(B)* GFP and *(C)* hBEST1-transfected HEK293T representative whole-cell current time courses at −100 mV and +100 mV during 1 min applications of external 30 mM GABA from a train of voltage ramps using the protocol shown in panel A. Cells were washed for 1.5 min with standard external solution between GABA applications. Blue circles represent mean current at −100 mV (bottom) or +100 mV (top) during the 250 ms steps at these voltages before and after the voltage ramp. Pink horizontal bars above the time course represent GABA application duration. Zero current level is denoted by the dotted black line. For time courses, GFP cells n = 3, hBEST1 cells n = 7. *(D)* Representative I-V relationships for the same hBEST1-transfected cell in the absence (black) and presence (pink) of external 30 mM GABA. Data are derived from the voltage protocols applied in *(C)*. “ctrl” indicates hBEST1 current without GABA. *(E)* Reversal potentials for hBEST1 currents in the presence and absence of external 30 mM GABA are plotted for different hBEST1-transfected cells from data as shown in *(D)* (n=7). Reversal potential measurement for hBEST1 without GABA (ctrl) was 0.8 ± 0.8 mV and with 30 mM GABA was 1.9 ± 1.2 mV. Mean is depicted by the black horizontal bar and error bars represent SEM. *(F)* GABA dose response. Representative hBEST1 whole-cell current time course at +100 mV using the protocol in *(A)* in the presence of various external GABA concentrations (from 300 μM to 50 mM), which were applied for 30 s, followed by washout for 1 min with external solution without GABA. 10 mM GABA was applied between each test concentration of GABA to control for possible rundown. *(G)* Dose response curve. Data from *(E)* in the presence of each GABA concentration were normalized to the subsequent current observed with 10 mM GABA. Data from all GABA titration recordings are shown as dots (n = 8); mean is depicted by black horizontal bars, with error bars representing SEM. Some error bars are smaller than symbols. Individual measurements are plotted as pink circles. *(H)* Representative hBEST1 whole-cell current time course with 30 mM GABA and 30 mM glycine with the same conditions as in *(B)* and *(C)*, n = 12. All recordings use a pipette solution containing 10 nM free Ca^2+^.

We next sought to assess whether the observed increase in current was due to an increase in Cl^-^ flow through the channel or due to GABA permeation itself. The GABA molecule (Fig. S1) would typically be considered a zwitterion at physiological pH indicating that it bears no formal charge; the pKas of the amine and carboxylate are estimated at 10 and 4, respectively (34). The possibility that GABA permeates through BEST1 and carries an ionic current in doing so (25) could potentially occur if specific interactions with the channel altered the pKa of either group to endow GABA with a charge at neutral pH. To address this possibility, we measured reversal potentials (the voltage at which the net current is zero) using voltage ramps, with symmetric Cl^-^, in the absence and presence of external GABA (Fig. 2*A* and *D-E*) (*Methods*). If an ion other than Cl^-^ permeates through the channel during GABA application, this would be detected by a shift in the reversal potential. We measured a reversal potential near 0 mV in both the absence and presence of GABA, indicating that GABA additions do not change the reversal potential (Fig. 2*D* and *E*). Therefore, the current observed upon GABA application is not due to GABA permeation through the channel. As observed in the fluorescence-based flux assay for chicken BEST1, the response of human BEST1 to GABA occurred in the millimolar range. Application of 300 µM GABA or less elicited no discernible increases in current, whereas application of 3 mM GABA or higher concentrations resulted in marked current increases (Fig. 2*F, G*). The response to GABA was not fully saturated at 50 mM, indicating a large dynamic range of response to GABA (Fig. 2*G*). To assess specificity and control for changes in osmotic pressure during high concentration additions of GABA, 30 mM glycine was used periodically in place of GABA. Glycine, which has a similar chemical structure to GABA, did not elicit increases in current (Fig. 2*H* and Fig. S1).

To further characterize the effect of GABA, we tested its effect using a higher (100 nM) concentration of Ca^2+^ in the pipette solution. In the absence of GABA, currents increase following whole-cell break-in and then decline over time, as expected, due to inactivation of the channel at this higher Ca^2+^ concentration (Fig. S2). Applications of GABA during the time course cause concomitant increases in current even during the inactivation process. Our whole-cell patch-clamp observations complement the flux assay results and indicate that extracellular GABA activates BEST1.

### Bilayer electrophysiological studies of chicken BEST1

To further assess the effect of GABA on the channel, we turned to bilayer electrophysiology, which we have previously used to investigate the function of purified chicken BEST1 (10, 28, 33). Bilayer electrophysiology is a reconstituted technique that allows exquisite control of the experimental conditions, including the protein, lipid composition, and aqueous solutions (35). One of its benefits is that the use of purified components allows one to interrogate whether an effect is direct or indirect. Additionally, the system facilitates changing the components of the aqueous solutions bathing either side of the membrane during the experiment. Using bilayer electrophysiology, we evaluated the effect of GABA on ionic currents through chicken BEST1. Using symmetric conditions, with identical (30 mM KCl) solutions on either side of the membrane, the addition of GABA markedly increases the current observed (Fig. 3*A*). A plot of the current increase as a function of GABA concentration can be fit with a standard activation curve (Fig 3*B*). Doing so indicates an EC_50_ of approximately 2 mM and a Hill coefficient of approximately 1.

**Fig 3.**
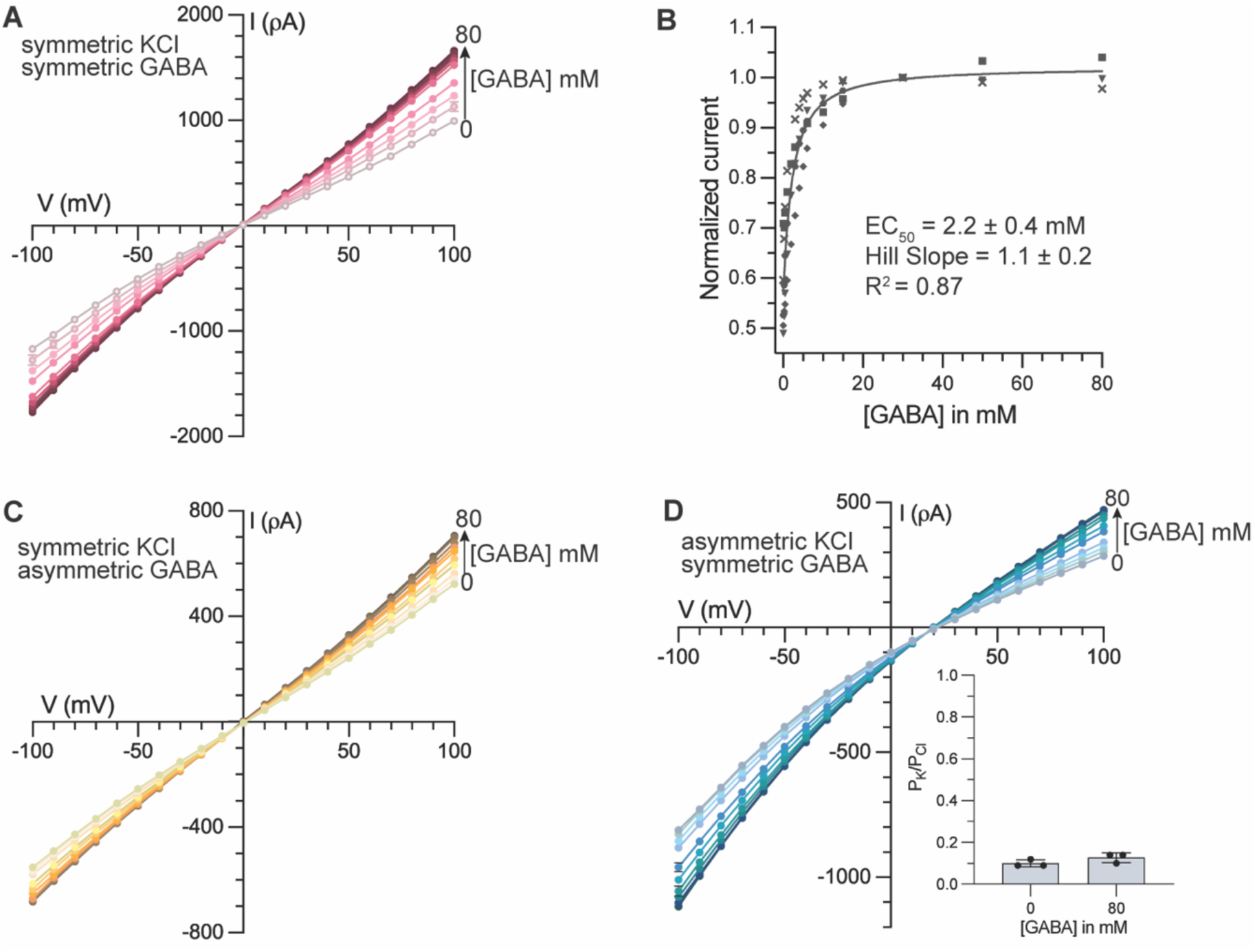
Bilayer electrophysiology shows that GABA potentiates Cl^-^ current through chicken BEST1. *(A)* Current-voltage (I-V) relationship indicates that GABA increases BEST1 currents. Symmetric conditions were used, such that both chambers contain 30 mM KCl. Increasing amounts of GABA were added to both chambers during the experiment. *(B)* GABA dose response curve. Currents observed at 100 mV at different GABA concentrations from *(A)* were plotted as fraction of current observed with 30 mM GABA at 100 mV. Data from five separate experiments, denoted by different symbols, were used to calculate half maximal concentration (EC_50_), Hill coefficient, and SE by fitting the data to a standard agonist activation function *(Materials and Methods)*. *(C)* Addition of GABA does not shift the reversal potential. The experiment was done in symmetric 30 mM KCl. Increasing amounts of GABA were added to only the cis chamber. *(D)* GABA does not change the channel’s selectivity for anions over cations. Asymmetric conditions were used, where the cis chamber contains 30 mM KCl and the trans chamber contains 10 mM KCl. Increasing amounts of GABA were added to both chambers. Permeability ratios (P_K_/P_Cl_) at 0 mM GABA and 80 mM GABA were calculated using the Goldman-Hodgkin-Katz equation; three separate experiments were used to calculate associated SE. Representative I-V graphs are shown in *(A, C, D)*; current traces for these are shown in (Fig. S3).

We next sought to revisit the possibility that GABA might be permeating through the channel in ionic form. To do so, GABA was added to only one side of the membrane, while other components of the buffers were held constant. The reversal potential would shift if the increase in current was due to GABA permeation in an ionic form. As expected, increases in current were observed with the addition of GABA to one side of the membrane (Fig. 3*C*). The extent of increase was lower than when GABA was added to both sides of the membrane, consistent with ion channels facing in both orientations as is typical in bilayer electrophysiology. I-V plots reveal that the reversal potential remains around 0 mV even when 80 mM GABA is added (Fig. 3*C*). The absence of a measurable change in reversal potential indicates that GABA does not detectably permeate through the channel in ionic form. The possibility that GABA may permeate through the channel in its zwitterionic form remains, but this cannot be detected with electrophysiology.

Finally, we sought to assess whether the GABA-dependent increase in current could be due to a change in the selectivity of the channel, perhaps by increasing the ability of the channel to permeate cations. Previous characterizations of the channel indicate that it preferentially permeates Cl^-^ in comparison to K^+^ (10). With 10 mM KCl on one side of the membrane and 30 mM KCl on the other, conditions we have previously used for such experiments, we observed a reversal potential of 21.9 ± 0.4 mV, indicating selectivity of the channel for Cl^-^ over K^+^ and a permeability ratio P_K_/P_Cl_ = 0.10 ± 0.01. If the effect of GABA is to change the channel’s relative permeability of Cl^-^ and K^+^, one would observe a change in the reversal potential. The addition of GABA reveals no change beyond the error in the measurement – the reversal potential is 20.6 ± 0.6 mV in the context of 80 mM GABA and P_K_/P_Cl_ = 0.13 ± 0.01 (Fig. 3*D*). We conclude that GABA does not significantly change the selectivity of the channel for Cl^-^ over K^+^. The data presented thus far indicate that GABA potentiates Cl^-^ currents by directly activating BEST1.

### Cryo-EM structures of human BEST1 in the absence of GABA

To contextualize the molecular basis for GABA potentiation, we sought to understand the conformational landscape of human BEST1 in the absence of GABA. Our single particle cryo-EM analysis of human BEST1 revealed two conformations of the channel (Fig. 4). By comparison with chicken BEST1 (9, 28), these two conformations represented: 1) an inactivated conformation where the neck is closed and the inactivation peptides are bound to their receptors, and 2) a conformation in which the neck of the channel is fully open, and the inactivation peptides are disordered. These structures are determined to 2.45 Å and 2.57 Å resolutions, respectively. We did not observe the ‘partially open’ conformation previously reported for human BEST1 (27). The inactivated structure is indistinguishable from that previously observed for human BEST1 (RMSD for Cα atoms = 0.3 Å) (27). The inactivation peptide, which has also been referred to as an auto inhibitory segment (27), binds to the cytosolic surface of the channel wherein the inactivation peptide (spanning amino acids 345 to 379) from each of the five BEST1 subunits wraps around two other subunits in belt-like fashion (Fig. 4*C*). In the inactivated structure, the neck of the channel adopts a closed conformation, as described previously (27, 28). Fifteen hydrophobic amino acids, three from each subunit (Ile76, Phe80, and Phe84), line the neck and form a constriction that is impervious to ions (Fig. 4*D*).

**Fig 4.**
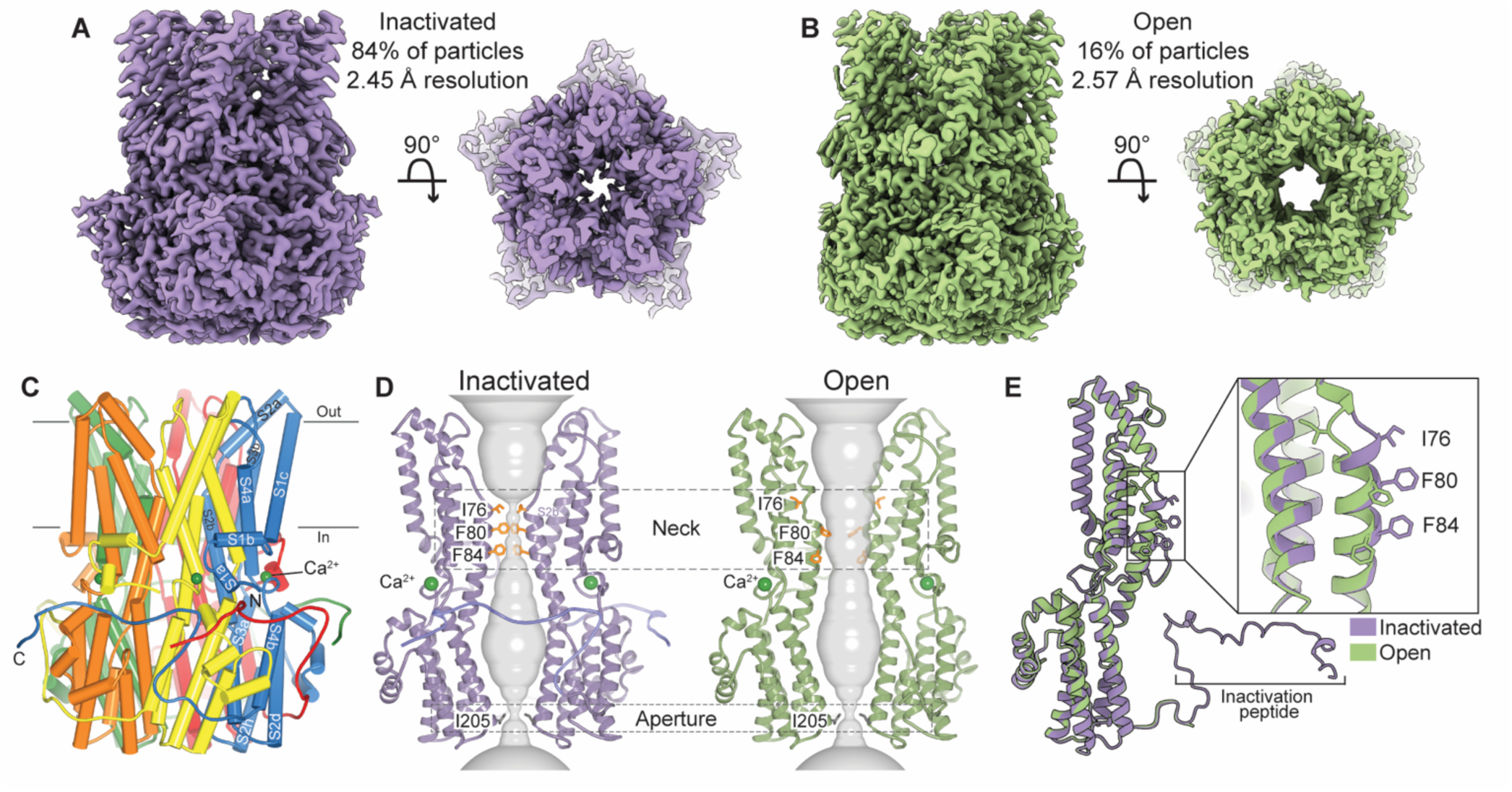
Cryo-EM structures of apo human BEST1. *(A-B)* Cryo-EM reconstructions of human BEST1 without GABA represent an inactivated state *(A)* and an open state *(B).* Left panels show views from the side; right panels show slices of the map from the extracellular side to highlight the widening of the neck in the open structure. *(C)* Overall architecture of inactivated human BEST1. α helices are depicted as cylinders and each subunit is colored differently. Approximate boundaries of a lipid membrane are indicated. *(D)* Cutaway views of the inactivated and open conformations of human BEST1. The ion conduction pore, show in grey, is depicted as the minimal radial distance from the center of the pore to the nearest van der Waals contact. The neck and aperture regions are indicated (dashed boxes). Two subunits are shown for clarity. *(E)* Superimposed individual subunits from the inactivated and open structures highlight the difference in conformation of the neck (boxed region). The inactivation peptide, labeled, is ordered in the inactivated structure but disordered in the open structure.

Our cryo-EM data set in the absence of GABA also revealed an open conformation of the channel (Fig. 4*B*). In this conformation, the inactivation peptide is disordered, and the neck of the channel is open. The phenylalanine residues in the neck that previously formed the constriction impervious to ions (Phe80 and Phe84) are now tucked away, and a large opening is formed that would be easily permeable to hydrated ions (Fig. 4*D*). The structure represents, to our knowledge, the first apo structure of BEST1 in an open conformation using a construct that contains the inactivation peptide. It is the same conformation as that obtained for both chicken and human BEST1 by removing the inactivation peptide (27, 28). These results confirm our biochemical, functional, and structural analyses of chicken BEST1 that indicated that the inactivation peptide dynamically interacts with its binding site on the cytosolic surface of the channel to allosterically control the conformation of the neck (28, 33). Both the open and the closed conformations of the neck are observed in the data set, indicating that these two states have similar energies. The mechanism by which the inactivation peptide biases the equilibrium is not fully understood. However, the open structure indicates that the neck can fully open even when the inactivation peptide is present.

Analysis of the human BEST1 data set also revealed dimers of complete channels wherein the cytosolic ends of two channels interact with one another, in a head-to-head manner (Fig. S5). The dimeric assembly, which we have not observed for chicken BEST1, may not be physiologically relevant; it would require that the two membranes in which the BEST1 channels reside would need to be separated by approximately 10 nm. Particles containing dimeric channels did not yield high resolution structures and therefore were excluded in the cryo-EM data processing. A comparison of the chicken and human BEST1 channels indicate the surface of the human channel is slightly more hydrophobic in the dimer interface, with a Gly198 in human BEST1 instead of Glu198 in chicken BEST1 (Fig. S5*B*).

### GABA shifts the equilibrium to favor the open state

Having obtained information on the conformational landscape of BEST1 in the absence of GABA, we sought to determine structures with GABA present. We found that inclusion of 30 mM GABA in the samples dramatically affected the landscape to favor the open conformation of both human and chicken BEST1. Without GABA, approximately 16% of particles of human BEST1 were observed in an open conformation and the remainder adopted an inactivated conformation. With GABA, approximately 66% of the particles adopted an open conformation (Fig. 5). The remainder adopted an intermediate state that is a structural hybrid between the open and inactivated states, described in more detail below. At the protein level, the open state conformation with GABA was structurally indistinguishable from the open conformation observed in the absence of GABA (Fig. 4*C*, Fig. 5*D*, RMSD for Cα atoms = 0.3 Å).

**Fig 5.**
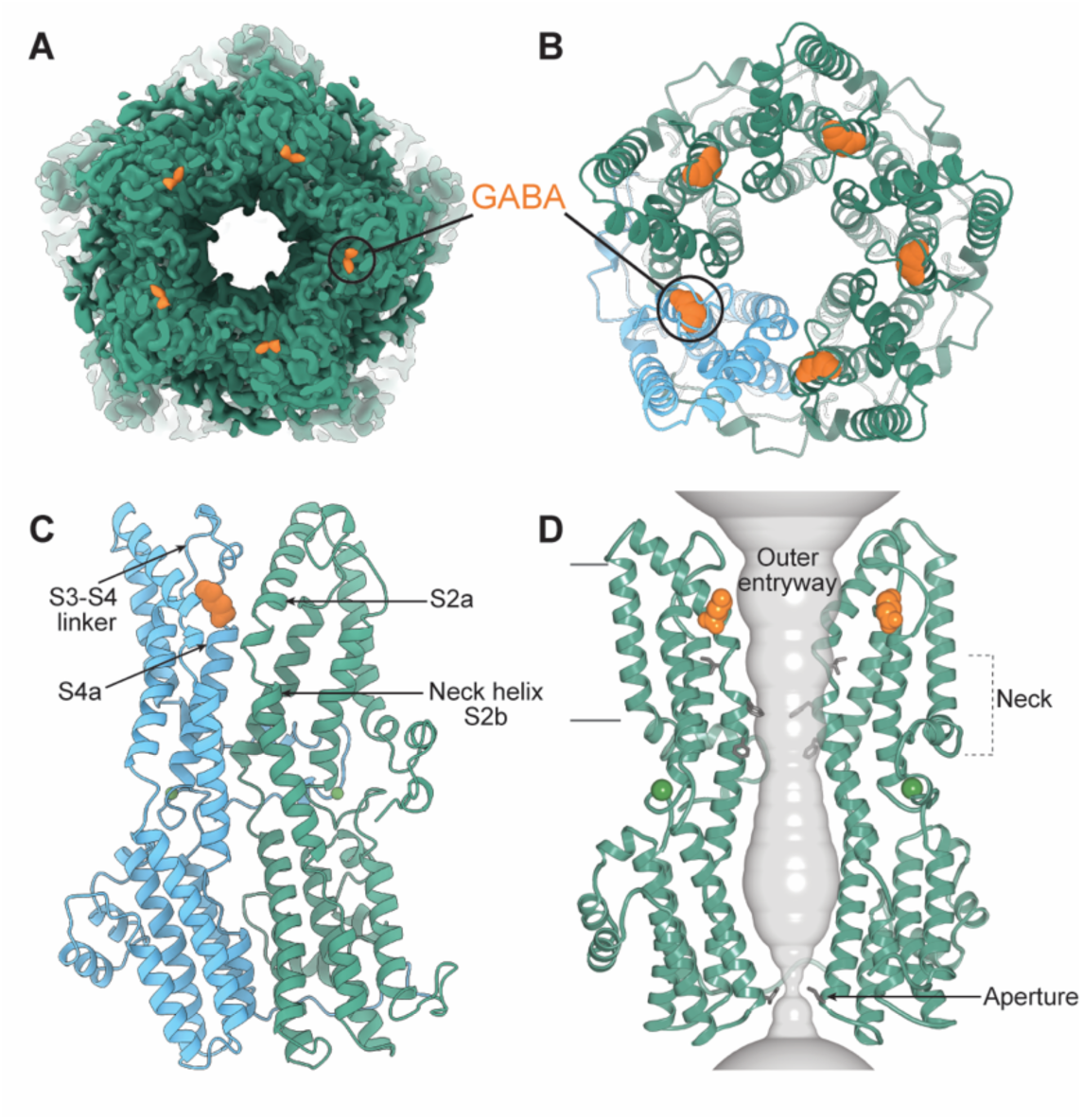
GABA-bound structure of human BEST1 in an open conformation. *(A)* 2.45 Å resolution cryo-EM map. Densities corresponding to GABA are colored orange. The view is a slice shown from the extracellular side, highlighting the open neck of the pore at the center. *(B)* Analogous view, showing a cartoon representation of the atomic model. GABA molecules are represented as orange spheres. One subunit is colored blue to highlight the binding of GABA at the interface between two subunits. *(C)* Side view of two adjacent subunits. GABA interacts with a portion of helix S2a from one subunit (green) and with the S3-S4 linker and amino-terminal end of the S4a helix from the adjacent subunit (blue). *(D)* Cutaway view of the GABA complex showing the pore (grey) and highlighting the binding of GABA adjacent to the pore within the outer entryway. The aperture adopts the same conformation in all structures of BEST1.

The structure of the open conformation of human BEST1 in the presence of GABA is determined to 2.45 Å resolution. At this resolution, the side chains of amino acids are well defined (Fig. S12). Non-protein densities consistent with GABA molecules are located at five symmetrical sites within the extracellularly-exposed ‘outer entryway’ of the pore (Fig 5). The sites are found at each of the five subunit-subunit interfaces (Fig. 5*B*).

We also determined structures of chicken BEST1 in the presence of GABA. Initially, we pursued structures of chicken BEST1 using a construct spanning amino acids 1-405, which contains the inactivation peptide. In the absence of GABA, this construct previously yielded a homogenous population of particles that represented the inactivated conformation of the channel (28). Cryo-EM analysis in the presence of 30 mM GABA also yielded the inactivated conformation of the channel; the neck of the pore was closed, the inactivation peptide was bound to its receptor, and no density could be ascribed to GABA.

Next, we used the construct of chicken BEST1 spanning amino acids 1-345 that lacks the inactivation peptide (28). Previous cryo-EM analysis of this construct yielded two conformations of the channel: a Ca^2+^-bound open conformation in which the neck was open, which represented ∼14% of the particles, and a Ca^2+^-bound closed conformation in which the neck was closed, representing the remaining majority. Strikingly, in the presence of GABA, the cryo-EM analysis yielded only an open conformation of the channel (Fig. S9). We were unable to discern a closed conformation from the dataset, despite this state representing the predominant conformation without GABA. The open structure with GABA is determined to 1.95 Å resolution (Fig. S9 and S10). The high resolution was useful for visualizing GABA, numerous ordered water molecules, and the conformations of amino acid side chains (Fig. 6*B* and S13). As for human BEST1, the GABA-bound open conformation of the protein is indistinguishable from the open conformation without GABA (RMSD for Cα atoms = 0.4 Å). Well-defined density for GABA was observed in the same location as for human BEST1 (Fig. 6*A* and *B*). We conclude that the presence of GABA alters the conformational landscape to favor the open conformation for both human and chicken BEST1.

**Fig 6.**
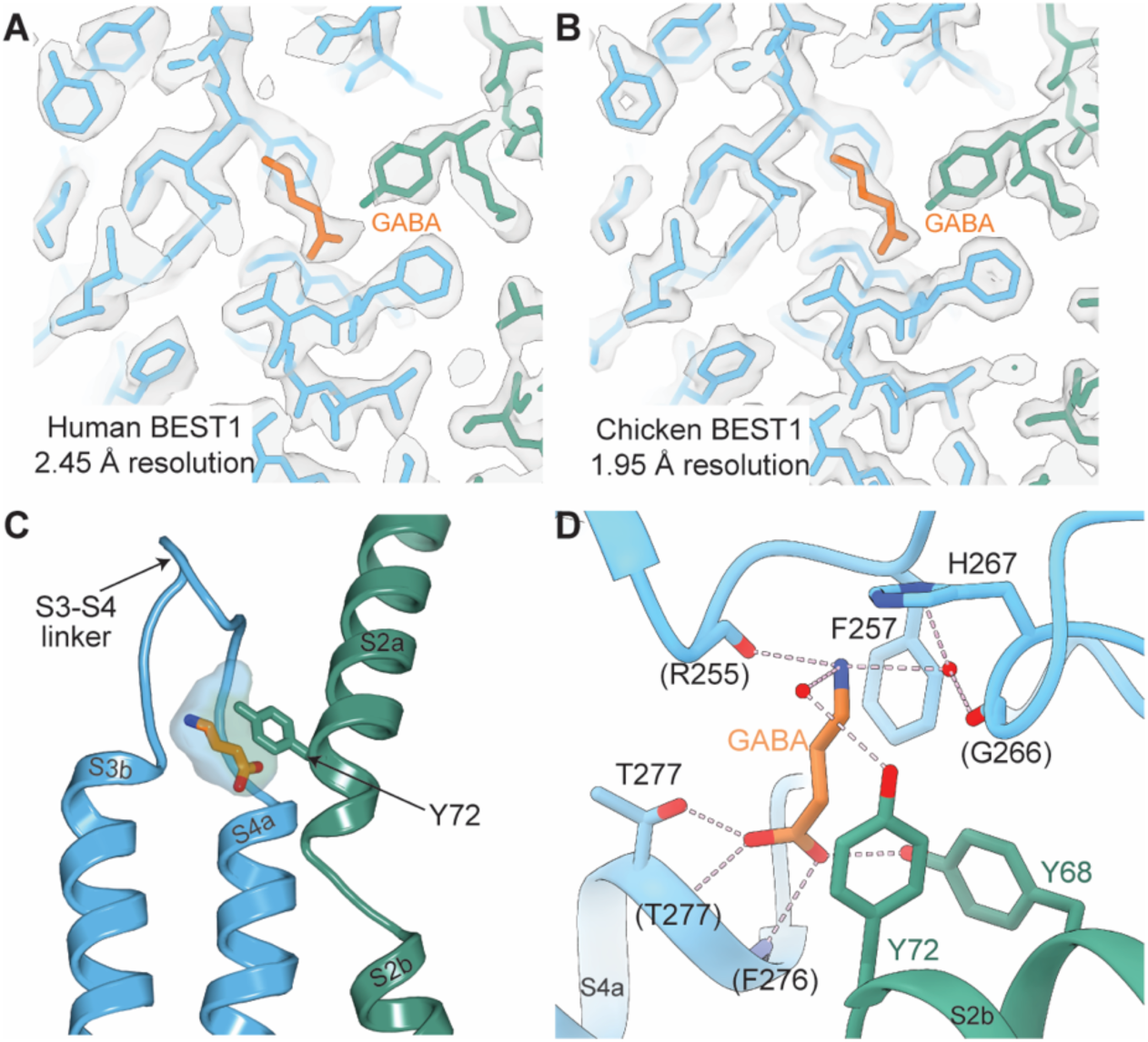
GABA binding site. One of the five identical binding sites for GABA is shown in each panel, with one subunit colored blue and the adjacent subunit green. *(A-B)* Cryo-EM densities from structures of the GABA-bound open conformations of human *(A)* and chicken *(B)* BEST1. Density is shown as a semitransparent surface. The atomic models are shown as sticks, with GABA colored orange. *(C)* Depiction of the GABA binding pocket, showing the cavity formed by the protein as a semitransparent surface. *(D)* Detailed interactions with GABA. The structure of human BEST1 with GABA is shown (the binding site in chicken BEST1 is analogous, Fig. S14). The atomic model is drawn as cartoons and sticks. Hydrogen bonds are depicted as dashed lines. Two ordered water molecules are shown as red spheres. Protein backbone atoms that form hydrogen bonds with GABA are designated by parentheses. Oxygen atoms are red; nitrogen atoms are dark blue.

### The GABA binding site

Five GABA binding sites are identified, according to the pentameric assembly of the channel. These sites are flanked by amino acids conserved between human and chicken BEST1 (Fig. 6 and Fig S14*C*). For both chicken and human BEST1, particles that represented the open conformation uniformly had GABA bound – we were unable to isolate classes of open channel particles that did not contain GABA, which is consistent with a high occupancy for GABA at these sites. Each of the five identical sites is formed at an interface between adjacent subunits. These sites had previously been identified as Cl^-^/Br^-^ binding sites from X-ray crystallographic analysis of chicken BEST1 in an inactivated conformation (9). GABA binds in a pocket on the sides of the wide ‘outer entryway’ of the pore that is open to the extracellular solution (Fig. 5*D*). The base of the pocket is formed by the amino terminal end of helix S4a and the S3-S4 linker (Fig. 6*C*). The pocket is ‘capped’ by Tyr72 and surrounding amino acids from the adjacent subunit. GABA makes extensive hydrogen bonding and van der Waals contacts with the protein. The carboxylate of GABA forms hydrogen bonds with two backbone nitrogen atoms at the amino terminal end of helix S4a (at Phe276 and Thr277), with Thr277, and with Tyr68 from the adjacent subunit (Fig. 6*D*). The amino end of GABA participates in a cation-pi interaction with Phe257, and it forms hydrogen bonds with the backbone carbonyl of Arg255 and two ordered water molecules, which are in turn coordinated by amino acids Tyr72 and His267. Tyr72 has extensive van der Waals contacts with the aliphatic portion of GABA. The aromatic ring of Tyr72 forms a thin separation between the pocket and the bulk solution of the outer entryway (Fig. 6*C*). In the GABA-bound structures, density for Tyr72 is well ordered, whereas density for this amino acid is weaker in the open conformation without GABA, suggesting that Tyr72 has increased mobility in the absence of GABA (Fig. S11 and S12). A dynamic nature of Tyr72 may facilitate access to the binding site.

### An intermediate structure

The single particle analysis of human BEST1 in the presence of GABA also revealed an intermediate conformation (Fig. 7*B* and Fig. S7). In this structure, GABA is bound at two of the five sites – in a GCGCC manner (G indicating GABA bound and C indicating unoccupied). We were unable to identify other structures with different configurations of GABA binding from the dataset. In the observed intermediate, the neck of the channel adopts a narrow configuration but with slight changes to the positioning of the neck-lining phenylalanine residues (Phe80 and Phe84) relative to the closed conformation without GABA (Fig. 7 *D*, *E* and Fig S7*B*). In the sites with GABA bound, amino acids Tyr72-Pro77 adopt the configuration observed in the fully open channel, and Tyr72 contacts GABA. In the unoccupied sites, Tyr72-Pro77 adopt their closed conformation. We refer to amino acids Tyr72-Pro77 as the ‘GABA switch’ because its conformation corresponds with the presence or absence of GABA.

**Fig 7.**
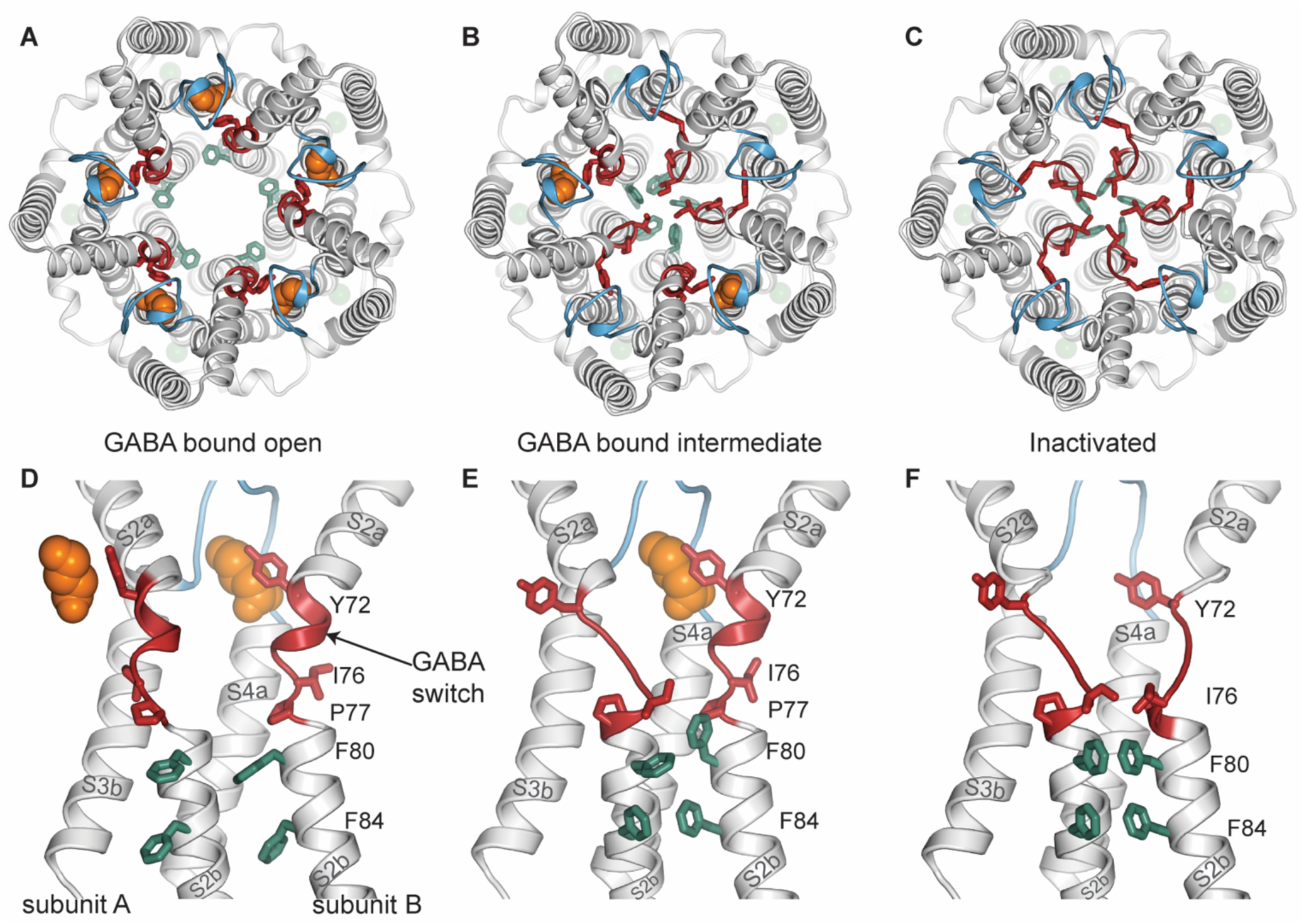
The GABA switch allosterically controls gating of the neck. *(A-C)* Comparison of cryo-EM structures of human BEST1 in the GABA-bound open conformation *(A)*, the GABA-bound intermediate conformation *(B)*, and the inactivated conformation in the absence of GABA *(C)*. The structures are depicted as cartoons, with certain amino acids drawn as sticks. The view is from the extracellular side, to highlight the dimensions of the neck. GABA molecules are depicted in sphere representation (orange). The S3-S4 linker is blue, the GABA switch (Tyr72-Pro77) is red, and Phe80 and Phe84 of the neck are green. Two GABA molecules are bound in the intermediate structure, as shown. Ca^2+^ ions are shown as faded green spheres. *(D-F)* close-up view of the GABA binding site, the GABA switch, and the neck from the structures. In each panel, the GABA switch and neck helix (S2b) from two adjacent subunits (labeled A and B) are shown, but only one S3b-S4a region is drawn (from subunit A). In the open conformation *(D)*, GABA is bound at all five sites, and the GABA switches and neck helices uniformly adopt their open conformations. In the inactivated conformation *(F)*, the neck and GABA switch regions adopt their closed conformations. The intermediate *(E)* is a structural hybrid. The GABA switch to which GABA binds (subunit B) adopts a GABA-bound conformation. The other GABA switch (subunit A) adopts a closed conformation. The neck of the intermediate is narrow *(B)* and similar to the inactivated state.

### The GABA switch and the mechanism of GABA activation

As described earlier from our studies of chicken BEST1 (28), opening of the neck involves a concertina of rotamer side chain rearrangements that allow the neck-lining phenylalanine residues (Phe 80 and Phe 84) to swing away from the central axis of the pore and thereby create a large opening within the neck (Fig. 7 *A* and *C*). The region of the polypeptide identified here as the GABA switch (Tyr72-Pro77) undergoes dramatic changes in its secondary structure between the open and closed conformations of the pore – these represent the largest backbone conformational changes within the entire polypeptide (Fig. 7 *D, E* and *F*). Tyr72 and Ile76, both part of the GABA switch, transition between components of α - helices to having extended secondary structures (Fig. 7 *D* and *F*). As a result, opening of the neck is accompanied by a shortening of the neck-lining S2b helix and a lengthening of S2a. Ile76, which is part of the neck helix (S2b) in the closed state, moves by approximately 9 Å (measured at its Cα atom) in the open state. Ile76 is no longer a part of the neck in the open state – rather a break in the neck helix occurs at Pro77 and the side chain of Ile76 binds in a hydrophobic cleft, formed by amino acids Phe247, Phe276, Leu279 and Phe283, on the side of the pore (Fig. S15*A*). Among the amino acids that contact GABA directly, Tyr72 undergoes the largest conformational change between the closed and open conformations, moving by approximately 4 Å (Fig. 7 *D* and *F*, Fig. S15*B*). In the opening transition, Tyr72 moves from being part of the linker between S2a and S2b to being a component of the S2a helix. The repositioning of the GABA switch creates the binding pocket for GABA. The ligand would be unable to bind to the closed conformation of the channel due to steric clashes, primarily with Tyr72 (Fig. 7 *D, E,* and *F*).

In comparison with the closed conformation, other small changes occur to accommodate GABA binding. Tyr68 moves by approximately 1.5 Å to form a hydrogen bond with the carboxylate of GABA (Fig S15*B*). A slight general expansion of the entire extracellular entryway also accompanies opening; for example, the amino end of helix S4a, with which GABA interacts, is positioned approximately 1 Å further from the center of the pore in the open conformation than in the closed conformation. Other changes are necessary to accommodate movement of the region of the GABA switch that does not interact directly with GABA – these include rotamer conformational changes of Gln280, Phe282, and Phe283 on the adjacent subunit to make room for Leu75, Ile76, and Pro77, as we have shown previously from comparisons of open and closed structures without GABA (Fig. S15*A*) (28).

Inspection of the structures engenders a mechanism of GABA-dependent activation (Fig. 7). GABA binding stabilizes the open conformation of the channel by stabilizing an open conformation of the GABA switch – when GABA is bound, the GABA switch is inhibited from adopting the conformation observed in the closed state. The GABA switch is a direct component of the neck of the channel in its closed conformation. Movement of the GABA switch is necessary for opening of the neck and for the binding of GABA. However, the intermediate structure indicates that GABA can bind to two of the five sites without opening of the neck. Therefore, there is a degree of flexible coupling, a ‘molecular clutch’, between the GABA switch and the neck of the channel. Taken together the data indicate that GABA stabilizes the open conformation of the neck through the conformational change in the GABA switch, and that multiple GABA binding events further favor channel opening.

## Discussion

### General summary

We have shown here that GABA, the chief inhibitory neurotransmitter in the central nervous system, activates the BEST1 chloride channel. Functional and structural studies of chicken and human BEST1 indicate that GABA activation is direct. There are five GABA binding sites on the channel, according to its pentameric architecture. The binding sites are located at interfaces between adjacent subunits and are exposed to the extracellular solution. Whole-cell patch-clamp electrophysiology indicates that the extracellular application of GABA activates the channel in a reversable manner. Cryo-EM studies reveal that GABA binding stabilizes the channel in a conductive conformation wherein the neck of the channel, which functions as its gate, is open. In each GABA site, the amino terminal end of helix S4a and surrounding residues from the same subunit form the base of the binding pocket, which retains a relatively fixed conformation between the open and closed states of the channel. The GABA switch from the adjacent subunit forms the cap of the binding pocket and undergoes substantial conformational changes between the open and closed states. Reorganization of the GABA switch is necessary to form the site. The GABA switch is directly connected to the S2b helix which forms the walls of the neck of the channel. The GABA binding site is thereby directly connected to the gating apparatus of the channel. Comparison among the structures indicates that GABA dramatically increases the percentage of particles that adopt the open confirmation. The data suggest that GABA binding thereby activates the channel by increasing the probability that the neck adopts an open conformation.

### GABA does not alter the ion selectivity of BEST1

The region(s) of the channel responsible for its selectivity for anions over cations are not known. Mutations of the neck and the aperture have been shown, for example, to have no effect on anion vs cation selectivity (10). Our structural data indicate that GABA binds at a site previously identified as a Cl^-^/Br^-^ binding site. We had hypothesized that the ability of this site to bind anions might contribute to the channel’s selectivity for anions (9). A structure-function analysis for this site is difficult to obtain because Cl^-^ is coordinated by backbone nitrogen atoms, an interaction that is inaccessible to mutation. The displacement of Cl^-^ by GABA allowed us to interrogate the role of this site in anion selectivity. The electrophysiology data presented here show that high (80 mM) concentrations of GABA that maximally activate chicken BEST1 do not detectably alter the selectivity of the channel for anions over cations. We conclude that the ability of this site to bind Cl^-^ is not responsible for the channel’s selectivity for anions over cations – this aspect of the channel’s function remains a mystery but could be partially due to a Cl^-^ site observed in the inner cavity of the channel (9).

### On GABA permeation

The electrophysiological data presented here indicate that GABA does not permeate through BEST1 in an ionic form to an appreciable degree at neutral pH, as would be expected due to its zwitterionic form. Our experiments do not address the possibility that GABA can permeate through the channel in zwitterionic form, as has been suggested (25), because electrophysiology cannot detect the permeation of uncharged species.

### Similarities with the GABA_A_ receptor

The GABA binding site of BEST1 has similarities to the GABA binding site in the GABA_A_ receptor (36) (Fig S14). As in BEST1, GABA binds to the GABA_A_ receptor at an interface between adjacent subunits. Hydrogen bonds are formed with both the carboxyl and amino ends of the GABA molecule in the GABA_A_ receptor. In GABA_A_, a tyrosine residue (Tyr205) packs against GABA, forming van der Waals contacts with much of the ligand, in a similar manner that Tyr72 does in BEST1. The carboxylate of GABA in the GABA_A_ receptor also forms hydrogen bonds with nitrogen atoms – in that case with an arginine side chain (Arg100) instead of the backbone nitrogen atoms observed in BEST1. Like BEST1, the GABA_A_ receptor is a pentameric chloride channel. However, the channels have no similarities in their overall structures, and the mechanisms of GABA-dependent activation are distinct. For instance, GABA binding in the GABA_A_ receptor occurs in this channel’s large extracellular ligand binding domain rather than within the transmembrane region as in BEST1.

### Concluding remarks

Many classes of cation channels share a common pore architecture and gating mechanism even though they are selective for different cations (e.g. Na^+^, K^+^, Ca^2+^) and are controlled by different stimuli, such as ligands or voltage (37). Different classes of anion channels, on the other hand, have proven to be structurally very distinct. Each class of anion channel discovered seems different than the rest with respect to structure and gating mechanism. For example, the two known types of Ca^2+^-activated chloride channels, bestrophin channels, and TMEM16A-type channels, could hardly be more different from one another from structural or mechanistic perspectives (9, 38). Bestrophin channels are pentamers with a single ion conduction pore. They are gated by Ca^2+^ binding to sites in the cytosolic domain of the channel. TMEM16A channels are dimers with two ion conduction pathways, each of which runs not down the middle of the protein but on its periphery and is exposed to the lipid membrane itself (39–41). In TMEM16A channels, Ca^2+^ activation occurs by binding within the membrane spanning region. Structural and electrophysiological studies of BEST1 have been fruitful in elucidating the unexpected mechanisms of this channel. We have found that the neck is both the Ca^2+^-dependent activation gate and the inactivation gate: Ca^2+^ binding favors opening of the neck and inactivation peptide binding favors its closing (10, 28, 33). In this study, we show that BEST1 is also activated by extracellular GABA and that GABA controls the same gate – the neck. Most other ligand-gated ion channels have ligand binding sites located on ligand binding domains outside of their pores. BEST1 is one of the few channels in which protein conformational changes that underlie gating are controlled by a ligand that binds at the periphery of its pore.

Sequence analysis and a structure of human BEST2, show that the site in which GABA binds BEST1 is conserved in BEST1-3 (Fig S14*C* and (27)). This sequence and structural conservation suggest that BEST1-3 may also be activated by GABA, but this remains to be tested experimentally. The site is similar in BEST4 in that all amino acids that coordinate GABA are conserved except for Tyr72, which is replaced by a serine residue. The extensive contacts that Tyr72 makes with GABA in BEST1 suggest that BEST4 may have diminished response to GABA.

The studies presented here show that extracellular GABA activates the BEST1 chloride channel. During the preparation of this work for publication, another study presented electrophysiological and cryo-EM studies that demonstrated activation of human BEST1 by GABA (29). Our work expands and corroborates this finding. A notable difference is the reported response range for GABA – we observe a response in the millimolar range, whereas the other study suggested a response in the ∼400 nM range. The reasons for this difference are unclear. We find that the channel displays specificity for GABA among the naturally occurring molecules that were tested, but it remains possible that other endogenous ligand(s) could activate BEST1. The mechanism of GABA-activation is to dilate the neck of the channel. The neck also serves as the gate for Ca^2+^-dependent activation and for inactivation, both of which processes are controlled by cytosolic inputs. Astonishingly, three inputs from two sides of the membrane impinge on one structural element to control the flow of ions through the channel (Fig. 8). The identification that BEST1 is activated by GABA will hopefully inspire new efforts to further probe the biological functions of the broadly expressed bestrophin family of channels.

**Fig 8.**
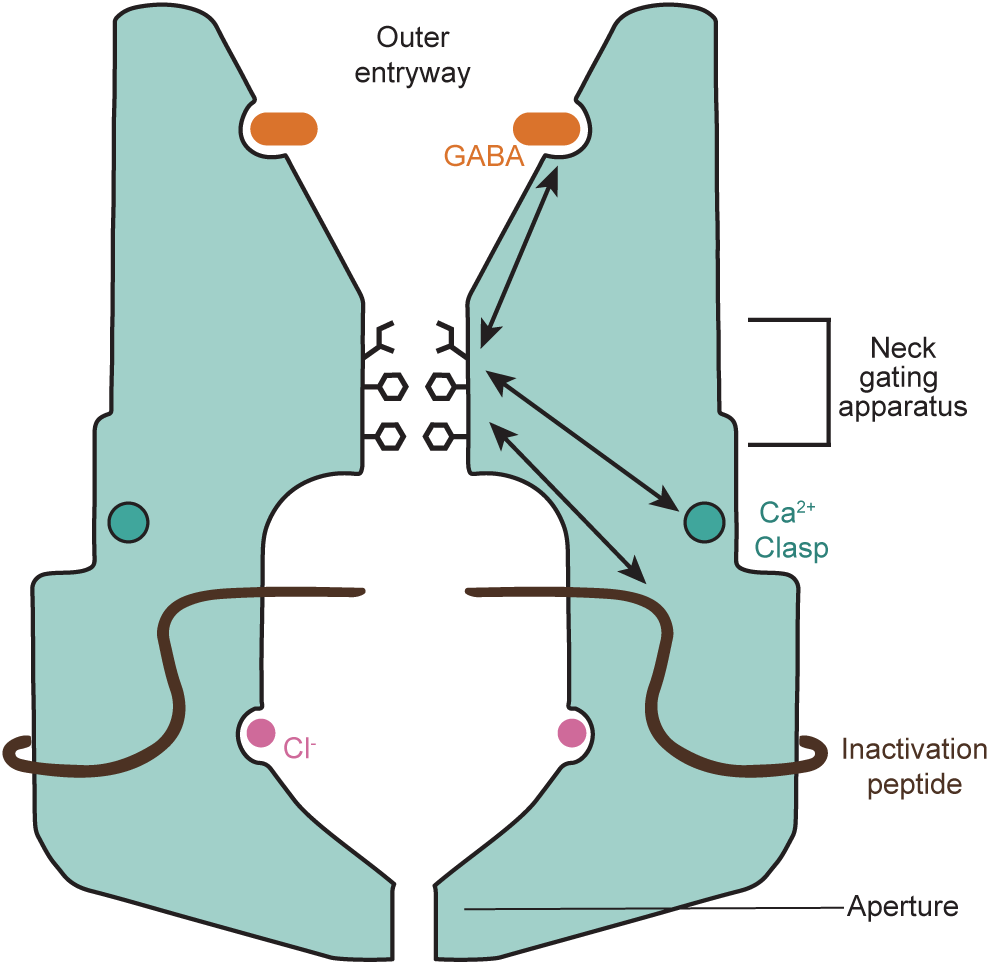
Gating. A schematic of the channel is shown. The neck serves as the gate of the channel. It is allosterically controlled by three inputs (arrows): Ca^2+^ binding to the cytosolic Ca^2+^ clasp sensor, binding of the inactivation peptide to a cytosolic receptor on the channel, and GABA binding within the outer entryway of the pore. The aperture does not function as a gate, but rather acts as a size selective filter to govern the size of anions that can flow through the channel.

## Materials and Methods

### Protein expression and purification

Chicken BEST1 constructs (spanning amino acids 1-405 and 1-345) were expressed in *Pichia pastoris* and purified for reconstitution into liposomes and cryo-EM sample preparation as previously described (9, 28). For chicken BEST1 reconstitution into liposomes, the size exclusion chromatography (SEC) buffer was: 20 mM Tris-HCl, pH 7.6, 150 mM NaCl, and 3 M n-decyl-β-D-maltopyranoside (DM, Anatrace). A Superose 6 Increase 10/300 GL column (GE Healthcare) was used at 4°C for all SEC purifications. For cryo-EM analysis, chicken BEST1 was purified using SEC in the buffer: 20 mM Tris, pH 7.6, 50 mM NaCl, 1 mM n-dodecyl-β-D-maltoside (DDM, Anatrace) and 0.1 mM cholesteryl hemisuccinate tris salt (CHS, Anatrace). After SEC, the sample (∼1 mL) was concentrated to 7 mg/mL, using a100-kDa MWCO concentrator (Amicon Ultra-2), and used immediately for cryo-EM grid preparation.

The human BEST1 protein (spanning amino acids 1-398) was expressed in *Pichia pastoris* using the pPICZ vector (Invitrogen) with a C-terminal green fluorescent protein (GFP) tag, separated by a linker for cleavage by PreScission protease. All steps for purification were done at 4°C. For reconstitution into liposomes, human BEST1 was extracted from 10 g of lysed cells using 1.75 g DDM for an hour in 150 mL lysis buffer [50 mM Tris-HCl, pH 7.5, 150 mM NaCl, 0.1 mg/mL DNase (Sigma-Aldrich), 1:600 Protease inhibitor cocktail set III (Sigma Aldrich), 0.5 mM AEBSF (Gold Biotechnology), 1:1000 Aproptonin (Sigma-Aldrich),1.5 μg/mL Leupeptin (Sigma-Aldrich), 1.5 μg/mL Pepstatin (Sigma-Aldrich) and 0.1 mg/mL Soybean Trypsin inhibitor (Sigma-Aldrich)]. The lysate was then centrifuged at 45,000 g for 1 hour at 4°C and the supernatant was filtered using a 0.22 μm filter (Millipore). The filtered sample was incubated with 3 mL of CNBr activated Sepharose Fast Flow resin (GE Healthcare) beads coupled to GFP nanobody (42) for an hour with gentle agitation. The beads were then applied to a column and washed with 70 mL of purification buffer containing 20 Tris-HCl, pH 7.6, 150 mM NaCl, 3 mM DDM, 0.15 mM CHS and 0.06 mg/mL of 3:1:1 (wt/wt/wt) of POPC:POPE:POPG (Avanti). The beads were then incubated with 4 mL of purification buffer containing 0.2 mg of PreScission protease and 1 mM dithiothreitol (DTT) for three hours at 4°C. Human BEST1 was eluted from the beads and was concentrated to approximately 500 ml using 100-kDa MWCO concentrator (Amicon Ultra-15). The concentrated sample was further purified using SEC in buffer containing 20 mM Tris-HCl, pH 7.6, 150 mM NaCl, 10 mM CaCl_2_, 1 mM DDM, 0.1 mM CHS and 0.06 mg/mL of POPC:POPE:POPG. The sample (2 mL) was concentrated to 9 mg/mL using a 100-kDa MWCO concentrator (Amicon Ultra-2). Concentrated sample was used immediately for reconstitution into liposomes.

For Cryo-EM analysis, human BEST1 was purified with the following modifications to the procedure described above. After binding to the GFP nanobody resin, the column was washed with 70 mL of purification buffer containing 20 mM Tris-HCl, pH 7.6, 150 mM NaCl, 10 mM CaCl_2_, and 0.5 mM glyco-diosgenin (GDN; Anatrace). The SEC buffer contained 40 mM HEPES, pH 7.6, 200 mM NaCl and 50 μM GDN to mimic conditions previously used for cryo-EM analysis of human BEST1 (27). After SEC, the sample (∼ 2 mL) was concentrated to 4 mg/mL using a 100-kDa MWCO concentrator (Amicon Ultra-2) and immediately used for cryo-EM grid preparation.

### Reconstitution into liposomes

Reconstitution of purified chicken BEST1 (1–405) into liposomes followed procedures previously described (9, 10). Briefly, for the fluorescence-based flux assay, a 3:1 (wt/wt) POPE:POPG lipid mixture (Avanti) was resuspended in reconstitution buffer: 10 mM HEPES, pH 7.0, 100 mM Na_2_SO_4_, 0.2 mM EGTA and 0.1 mM CaCl_2_. For planar lipid bilayer electrophysiology, the 3:1 (wt/wt) POPE:POPG lipid mixture was resuspended in 20 mM HEPES, pH 7.6, 450 mM NaCl, 0.2 mM EGTA and 0.19 mM CaCl_2_. Resuspended lipids (at 20 mg/mL) were sonicated for total of 6-10 minutes and then solubilized using 8% (wt/vol) of n-octyl-β-D-maltoside (OM; Anatrace). Purified chicken BEST1 and SEC buffer, as needed, were added to the solubilized lipids to yield protein:lipid (wt/wt) ratios of 1:100, 1:500, such that the final lipid concentration was 10 mg/mL. Empty vesicles (i.e. liposomes without purified BEST1) were also prepared in parallel using SEC buffer in place of purified protein. The samples were then incubated at room temperature for one hour without agitation. To remove detergent, the mixture was dialyzed using 8 kDa molecular weight cutoff dialysis tubing (Spectra/Por 7 Membrane Tubing; Fisher Scientific Catolog no. 086805A) in reconstitution buffer at 4°C for 5 days with daily 2 L buffer changes. Following dialysis, vesicles were sonicated briefly, aliquoted into 40 - 60 μL aliquots, flash frozen, and stored at −80°C until use.

The procedure for reconstitution of human BEST1 (1–398) into liposomes generally followed methods described for the MCU channel (43). A 3:1 (wt/wt) POPE:POPG lipid mixture was prepared for reconstitution and resuspended in reconstitution buffer (5 mM HEPES, pH 7.0, 100 mM K_2_SO_4_, and 200 μM CaCl_2_) to yield lipid concentration of 20 mg/mL. Lipids were sonicated for 6-10 minutes. 140 μg of human BEST1 purified in DDM (9 mg/mL) was added to 14 mg of lipids to yield a 1:100 protein:lipid (wt/wt) ratio and the mixture was incubated overnight at 4°C with end-over-end agitation. Empty liposomes were also prepared in parallel by adding SEC buffer in place of purified protein. 100 mM methyl-β cyclodextrin (MBCD; Sigma-Aldrich), dissolved in the reconstitution buffer, was added to the lipid-protein mixture so that the final concentration of MBCD was ∼1.2 mM. The sample was rotated end-over-end at 4°C. The same amount of MBCD was added to the sample every 8 hours, with a total of 4 additions. The vesicles were then pelleted by centrifugation at 194,000 g (TLA 100.3 fixed angle rotor) for 1 hour at 4°C. The supernatant was discarded, and the vesicle pellet was resuspended in reconstitution buffer so that the final lipid concentration was 10 mg/ml (based on full recovery of the initial lipids). Vesicles were sonicated briefly, aliquoted into 60 μL aliquots, flash frozen and stored in −80°C until use.

### Fluorescence based flux assay

The flux assay followed methods previously published (9). Briefly, liposomes were thawed rapidly (using a 37°C water bath), sonicated for 20 seconds, and then incubated at room temperature for 2 hours before use. For chicken BEST1, the flux assay buffer consisted of 10 mM HEPES, pH 7.0, 0.2 mM EGTA, 0.1 mM CaCl_2_, 2 μM 9-amino-6-chloro-2-methoxyacridine [ACMA; 2 mM stock in dimethyl sulfoxide (DMSO)], 0.5 mg/mL bovine serum albumin, and the indicated test compound (NaCl, GABA, etc.). Detailed buffer compositions for each flux assay experiment are given in Table S1. Na_2_SO_4_ was used to balance osmolality for test compounds as indicated (Table S1). Unless noted otherwise, all reagents were purchased from Sigma-Aldrich. Fluorescence was measured throughout the experiment every 30 sec. At the 1-minute mark, liposomes were mixed into the flux assay buffer to yield a 100-fold dilution (10 μL liposomes into 1 ml of flux assay buffer). At 2.5 minutes, 2 μM of the proton ionophore carbonyl cyanide m-chlorophenyl hydrazone (CCCP, from a 2 mM stock in DMSO) was added and mixed with a pipette. The flux assay with human BEST1 was completed with minor modifications. The flux assay buffer consisted of 5 mM HEPES, pH 7.0, 2 μM ACMA, 0.5 mg/mL bovine serum albumin, 200 μM CaCl_2_, and the indicated test compound (NaCl, GABA, etc.). Detailed buffer compositions are given in Table S1. Na_2_SO_4_ was used to balance osmolality for test compounds as indicated (Table S1). Data were collected on a SpectraMax M5 fluorometer (Molecular Devices) using SoftMax Pro 7 software. Fluorescence intensity measurements were collected every 30 seconds with excitation and emission wavelengths of 410 nm and 490 nm respectively. The data were normalized to the maximum fluorescence observed before adding CCCP. Data were analyzed and visualized using GraphPad Prism 10.

### Whole-cell patch-clamp electrophysiology

For expression in mammalian cells, human BEST1 cDNA (spanning amino acids 1-398) was cloned into the XhoI and EcoRI sites of a mammalian expression vector that includes a C-terminal Rho-1D4 antibody tag (44, 45). Sequences were confirmed by DNA sequencing (Azenta).

Human Embryonic Kidney 239T (HEK293T, ATCC cat # CRL-3216) cells were cultured in Dulbecco’s modified Eagle’s medium (DMEM, Gibco cat # 1995065) supplemented with 10% fetal bovine serum (FBS) (Gibco cat # A3840002) and 100 µg/mL of kanamycin. HEK293T cells between passage numbers 5-30 were used and passaged when cells were between 40-80% confluent. In preparation for patch-clamp experiments, cells were treated with 0.25% trypsin for 20-30 s, resuspended in warmed cell media, and then seeded on glass coverslips (Fisher Scientific, 22 x 22 mm No. 1 thickness, cat # 50-189-9778) at approximately 15% of the original confluency in 35 mm petri dishes. Transfections were done using the FuGENE6 Transfection Reagent (Promega, following manufacturer instructions) using a 3:1 ratio of FuGENE6 to DNA; 6 µL of FuGENE6 to 2 µg DNA. To facilitate identification of transfected cells, the hBEST1 expression vector was co-transfected with a vector containing cDNA encoding green fluorescent protein (GFP) (46) with a DNA ratio of 5:1, hBEST1 to GFP. Cells transfected with GFP alone also served as controls. Transfected cells were incubated for 16-48 hours at 37°C with 95% air and 5% CO_2_ before use in patch-clamp recordings.

Membrane currents were recorded from HEK293T cells using the whole-cell patch-clamp technique. When the whole-cell configuration was established, the desired extracellular solution was continuously perfused through a valve-controlled (Valvelink 8.2) gravity-fed perfusion system that used an 8-into-1 manifold (Automate Scientific) connected to a perfusion chamber (Warner Instruments, RC-26GLP). Membrane voltage was controlled using an Axopatch 200B patch-clamp amplifier (Axon Instruments) and currents were digitized using a Digidata 1550B and pCLAMP 11 software. Membrane currents were collected with a sampling rate of 10 kHz with a 2 kHz low-pass filter. Micropipettes were pulled with a P-1000 Next Generation Micropipette Puller (Sutter Instrument) using borosilicate glass with a filament (Sutter Instrument, cat # BF150-86-10HP). Typical patch pipette resistance ranged between 2-4 MΩ. Whole-cell recordings were performed starting from a holding voltage of −60 mV. Voltage ramps from −100 mV to +100 mV (over 500 ms, and flanked by 250 ms steps at −100 mV and +100 mV, respectively) were repeatedly applied every 2 s (Fig. 2*A*). Whole-cell current time courses depict the mean current values at each −100 mV and +100 mV step unless otherwise mentioned. Whole-cell patch-clamp recordings were excluded from analysis when: currents were too large (>5 nA), pipettes became obstructed and series resistance increased during the recording, or the cell was lost before the experiment could be completed.

Following previous characterization of BEST1 (32, 47), the standard extracellular solution contained 145 mM NaCl, 2 mM CaCl_2_, 1 mM MgCl_2_, 10 mM HEPES, and 15 mM D-glucose, adjusted to pH 7.4 with 1 M NaOH, with an osmolality of approximately 330 mmol/kg. The “zero” Ca^2+^ pipette solution contained 146 mM CsCl, 2 mM MgCl_2_, 5 mM EGTA, 10 mM HEPES, and 10 mM sucrose, adjusted to pH 7.3 with NMDG, with an osmolality of approximately 330 mmol/kg. A high Ca^2+^ (∼25 µM free Ca^2+^) pipette solution contained 146 mM CsCl, 2 mM MgCl_2_, 5 mM Ca-EGTA, 10 mM HEPES, and 10 mM sucrose, adjusted to pH 7.3 with NMDG, with an osmolality of about 330 mmol/kg. To achieve pipette solutions with the desired concentrations of free Ca^2+^, the “zero” Ca^2+^ and high Ca^2+^ pipette solutions were mixed according to free Ca^2+^ calculations performed with MaxChelator (48). A ground chamber contained the internal pipette solution and a ground electrode made with chlorided silver (Ag/AgCl) wire. The bath and ground chambers were connected by an agar bridge containing 3 M KCl. K^+^ was not used in recording solutions to prevent contamination from Kv channel currents. A 4 M GABA stock solution was made in deionized water and was used to make external solutions with the desired concentrations of GABA in the standard extracellular solution. A 1 M glycine stock made in deionized water, pH 8.0 with NaOH, was used to make an external solution containing 30 mM glycine. External solutions containing 30 mM GABA or 30 mM glycine had osmolalities of approximately 365 and 360 mmol/kg, respectively. Baseline current subtraction was not performed due to inactivation of hBEST1 in different intracellular calcium conditions. Electrophysiology data analysis and visualization were performed with Python (matplotlib) and GraphPad Prism 10.

### Planar lipid bilayer electrophysiology

Bilayer methods were based on those previously described (10). Liposomes containing chicken BEST1 (1–405) were thawed in 37°C water bath, sonicated for ∼10 seconds, and incubated at room temperature for 1 hour prior to use. The two chambers of the bilayer apparatus (Warner Instruments) were filled with 4 mL of bath solutions. Bath solutions contained 20 mM HEPES, pH 7.6, 0.21 mM EGTA, 0.19 mM CaCl_2_, and 10 mM KCl (trans chamber) or 30 mM KCl (cis chamber). Chlorided silver (Ag/AgCl) electrodes were submerged in 3 M KCl and connected to the bath solutions with salt bridges made with 2% agar (wt/vol) and 3 M KCl. Bath solutions were separated by a polystyrene partition with a 200 μm hole across which a bilayer was painted using POPE:POPG in n-decane [3:1 (wt/wt) ratio at 20 mg/mL]. Liposomes were added to the cis chamber 1 μL at a time until currents were observed, indicating that liposomes had fused with the bilayer. Each experiment began with recordings in asymmetrical (10/30 mM) KCl conditions to verify that the recorded currents displayed selectivity for Cl^-^ (from a reversal potential of ∼ 21 mV). For recordings in symmetric KCl conditions, after fusion of channels into the bilayer, the KCl concentration in the trans chamber was raised to 30 mM using 3 M KCl (while stirring).

For GABA activation experiments, KCl was made symmetric (cis/trans = 30/30 mM), and GABA (Millipore Sigma; 1, 2, or 4 M stock solutions in deionized water) was added to obtain the desired concentration (stirring for ∼1 minute). To measure GABA activation, currents at 100 mV were analyzed for each GABA concentration. To normalize between experiments using different bilayers (and containing different numbers of channels), the currents (which ranged from ∼140 pA to ∼900 pA at 100 mV) were normalized to the current observed when 30 mM GABA was present. To analyze the half maximal concentration (EC_50_) and Hill slope, the data were fit using the [agonist] vs. response – variable slope equation in GraphPad Prism 10.

For GABA permeation experiments, GABA was only added to the cis chamber under symmetric 30 mM KCl conditions to observe if the addition of GABA influenced the reversal potential. A representative I-V relationship is shown in Fig 3*C*; two analogous experiments are shown in Fig S3*G* and *H*. For experiments investigating whether GABA changes the selectivity of anions over cations, recordings were done in asymmetric KCl conditions (cis/trans = 30/10 mM), and GABA was added to both chambers. Relative permeabilities (P_K_/P_Cl_) were calculated from the measured reversal potential (*E_rev_*) and the Goldman Hodgkin Katz equation.

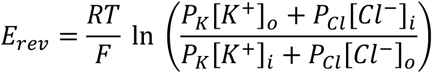

All bilayer electrophysiological recordings were made using a Warner Instruments planar lipid bilayer workstation. Currents were recorded using an Axopatch 200B amplifier (Axon Instruments), filtered at 1 kHz, and digitized at 5 kHz using the Clampex 10.4 program (Axon Instruments). Data were analyzed using Clampfit 10.4 (Axon Instruments) and GraphPad Prism 10. Currents from bilayers without channels are subtracted. All experiments were repeated at least three times, using different bilayers. Error bars represent SEM. The cis chamber is defined as the side to which vesicles are added. The trans chamber is defined as electrical ground.

### Cryo-EM sample preparation and data collection

All samples for cryo-EM were applied to glow discharged Quantifoil R 1.2/1.3 grids (Au 400; Electron Microscopy Sciences) and plunge frozen in liquid ethane using a Vitrobot Mark IV (FEI Thermo Scientific). For structural studies of chicken BEST1 with GABA, purified protein was concentrated to 7 mg/mL and filtered using a 0.22 μM spin filter. Immediately before the sample was applied to grids, 1 μL of GABA (300 mM stock) was added to 10 μL of protein sample to yield a 30 mM GABA concentration. The grids were plunge frozen using the Vitrobot (operated at 22°C, blot time of 3.5 seconds, wait time of 20 seconds, 0% blot force and 100% humidity). Micrographs were collected using a Titan Krios G4 microscope (Thermo Fisher) operated at 300kV using Falcon IV direct detector with a SelectrisX energy filter at the National Center for CryoEM Access and Training (NCCAT). Details of all datasets are summarized in Table S2. Similarly, purified human BEST1 was concentrated to 4.1 mg/mL and filtered using a 0.22 μm spin filter. For apo human BEST1 samples, the purified protein was applied to grids and plunge frozen using the Vitrobot (operated at 4°C, blot time of 3 seconds, wait time of 30 seconds, 0% blot force and 95% humidity). For studies with GABA, 30 mM GABA was added prior to grid preparation in the same manner as for chicken BEST1. Micrographs were collected using Titan Krios microscope (Thermo Fisher) operated at 300kV using Falcon IV direct detector SelectrisX energy filter (Thermo Fisher) at Memorial Sloan Kettering Cancer Center (MSKCC).

### Cryo-EM structure determination

Figures S4, S6 and S8 show cryo-EM data processing workflows. Data processing was done using Relion 3.1 (49) and cryoSPARC 4.5 (50). Data processing for human BEST1 without GABA proceeded as follows. Movies in EER format were fractionated into 45 fractions (up sampling factor of 1), and gain and motion-corrected using Relion 3.1 (49). These micrographs were imported into cryoSPARC 4.5 (50) for patch CTF estimation. Micrographs with a CTF fit better than 3.5 Å (6,415 micrographs) were selected for further processing. To obtain an initial model of human BEST1, 2000 micrographs were randomly selected.

Using these micrographs, particles were picked using blob picker and extracted (box size 384, bin factor 2). Three ab-intio reconstructions were generated and used for five rounds of heterogenous refinement. The resulting class that resembled BEST1 was used to create templates for particle picking, and particles were selected using the template picker tool in cryoSPARC. 5,338,248 particles were picked and extracted and binned by factor of 2. To remove junk particles, particles were sorted using multiple rounds of iterative 3D heterogenous refinement using the refined ab-initio model and decoy models (employing C1 symmetry). The resulting 1,357,422 particles were reextracted using a box size 512. Using 2D classification, the particles were classified into sets that contained monomer channels (pentamers) and dimers of channels (containing two pentamers in head-to-head assembly). Ab-initio dimer and monomer reconstructions were generated and used as inputs for heterogenous refinement of the particles along with decoy reconstructions. From this, 1,046,173 particles classified as monomers were subjected to further 3D classification using a solvent mask excluding the detergent micelle. This resulted in two conformations of human BEST1, inactivated and open, which were subsequently processed separately. Both particle sets were subjected to non-uniform refinement, followed by two rounds of Bayesian polishing in RELION 3.1, with heterogenous refinements in cryoSPARC 4.5 between polishing steps. Final particles (inactivated: 565,261, open: 107,143) were subjected to non-uniform refinement using C5 symmetry, with global CTF and defocus refinements, resulting in final maps of the inactivated and open conformations with 2.45 Å and 2.57 Å overall resolutions, respectively.

For human BEST1 with GABA, processing followed the same general procedure. Movies in EER format were fractionated into 45 fractions (up sampling factor of 1), and gain- and motion-corrected using Relion 3.1 (49). These micrographs were imported into cryoSPARC 4.5 (50) for patch CTF estimation. Micrographs with a CTF fit better than 3.5 Å (5,833 micrographs) were selected for further processing. 4,996,553 particles were picked and extracted from micrographs. To remove junk particles, particles were sorted using rounds of iterative heterogenous refinement using C1 symmetry. The resulting 962,724 particles were reextracted using a box size of 512. Dimer and monomer particles were separated as described above. 619,096 monomer particles were subjected to 3D classification, using a solvent mask excluding the detergent micelle. This yielded two discernable conformations, GABA-bound open (177,595 particles) and intermediate (163,534 particles). The volumes generated from 3D classification were used as input volumes, along with decoy models, for iterative heterogenous refinement. Iterative heterogenous refinement and 3D classification yielded two conformations, open (305,050 particles) and closed (180,153 particles). The two particle sets were subsequently processed separately. The particles in the open conformation class were subjected to non-uniform refinement, followed by two rounds of Bayesian polishing in RELION 3.1, with heterogenous refinements in cryoSPARC 4.5 in between polishing steps. The final particle set (295,666 particles) was subjected to non-uniform refinement in C5 symmetry, with global CTF refinement and defocus refinements resulting in a 2.45 Å overall resolution reconstruction of the open conformation. As the particle set for closed channels was still heterogenous, further processing was done to isolate different states. The particles were subjected to homogenous refinement with C5 symmetry in cryoSPARC 4.5, followed by C5 symmetry expansion. The symmetry expanded particle set was subjected to 3D classification, using 6 classes, a solvent mask excluding the detergent micelle, and a focused mask containing the neck region of BEST1 and the GABA binding site. 3D classification yielded multiple 3D reconstructions that contained two GABA molecules at the same positions within the pentamer (GCGCC configuration, i.e. at the interfaces between subunits A-B and C-D). The GABA switch regions at these positions also bore the same conformation in the classes. We were not able to obtain reconstructions with other distributions of GABA molecules or GABA switch conformations. Duplicate particles were removed, and the resulting 148,312 particles were subjected to local refinement in C1 symmetry to yield the intermediate reconstruction at 2.62 Å overall resolution.

Processing of data for chicken BEST1 (1–345) with GABA was similar. Movies in EER format were fractionated into 45 fractions (up sampling factor of 1), and motion-corrected using Relion 3.1 (49). Motion-corrected micrographs were imported into cryoSPARC 4.5 (50) for patch CTF estimation. Micrographs with a CTF fit better than 4.5 Å (1,224 micrographs) were selected for further processing. Following template picking, 987,499 particles were extracted using a box size 512 (binning by a factor of 2). The particles were subjected to iterative rounds of heterogenous refinement in C1 symmetry. Following unbinning, the particles were subjected to two rounds of 3D heterogenous refinement, yielding 75,954 particles in the open conformation. These particles were subjected to non-uniform refinement, followed by two rounds of Bayesian polishing in RELION 3.1, with heterogenous refinements in cryoSPARC 4.5 between polishing steps. The final particle set was subjected to non-uniform refinement in C5 symmetry, with global CTF refinement and defocus refinement, resulting in a reconstruction of the open map at 1.95 Å overall resolution. The dataset appears to have a single major conformation (open with GABA bound); multiple attempts were made to distill other conformations (e.g., closed, or those without GABA) by ab-initio reconstruction, 2D classification, 3D classification, or focused classification approaches. A relatively small fraction of particles from the initial set were used for the final map because this subset of particles yielded the highest resolution reconstruction and most well-defined density.

Atomic models were built in COOT (51) using previously determined structures of chicken and human BEST1 as references (PDB IDs: 4RDQ, 6N28, and 8D1I). All models were refined in real space using COOT and further refined using PHENIX (52). Structural figures were prepared using ChimeraX (53), PyMOL (pymol.org), and HOLE (54). Data collection and refinement statistics are shown in Table S2.

## Acknowledgements

We thank members of the S.B.L. laboratory and A. Accardi, O. Boudker, and M. Diver for helpful discussions; A. Miller for the pPICZ vector with a C-terminal GFP tag; M. Diver for sharing the patch-clamp rig; M. J. de la Cruz for assistance with cryo-EM at MSKCC; and the staff of the National Center for CryoEM Access and Training. This work was supported by NIH grant R35GM131921 (to S.B.L.), NIH core facilities grant to MSKCC (P30CA008748), NIGMS-funded Molecular Biophysics Training Program T32 predoctoral fellowship grant number 5T32GM132081 (to S.W.T.). A portion of the work was performed at the National Center for CryoEM Access and Training (NCCAT) and the Simons Electron Microscopy Center located at the New York Structural Biology Center, supported by the NIH Common Fund Transformative High Resolution Cryo-Electron Microscopy program (U24 GM129539, and NIGMS R24 GM154192) and by grants from the Simons Foundation (SF349247) and NY State Assembly.

## Data and materials availability

Atomic coordinates and maps of the apo human BEST1 (inactivated and open), GABA-bound human BEST1 (open and intermediate), and GABA-bound chicken BEST1 (open) have been deposited in the PDB (accession numbers: 9EGS, 9EGT, 9EGM, 9EGQ and 9EFZ, respectively) and EMDB (EMD-47996, EMD-47997, EMD-47991, EMD-47995 and EMD-47982, respectively). All data needed to evaluate the conclusions in the paper are present in the paper and/or the Supplementary Materials.

