## Supplemental Information for "The pentameric chloride channel BEST1 is activated by extracellular GABA"

**This PDF file includes:**

Figures S1 to S15
Tables S1 to S2

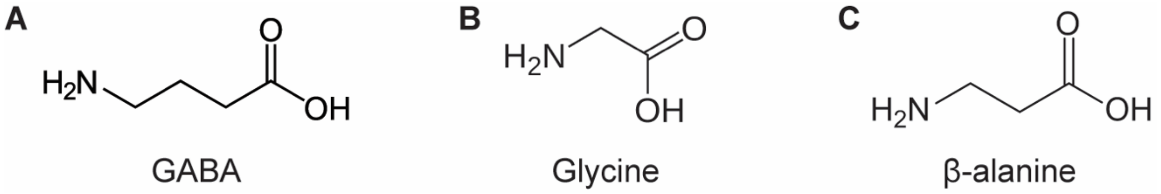


**Fig S1**. Chemical structures of GABA, Glycine, and β-alanine.

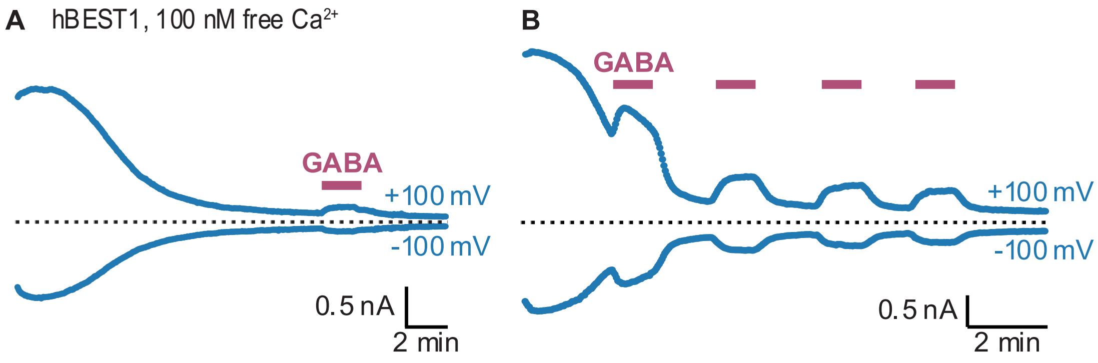


**Fig S2**. GABA and human BEST1 whole-cell current with intracellular 100 nM free $Ca^{2+}$. hBEST1 representative whole-cell current time courses at -100 mV and +100 mV from a train of voltage ramps (Fig. 2*A*), showing inactivation hBEST1 currents over time with 100 nM free $Ca^{2+}$ solution in the pipette. *(A)* Application of 30 mM GABA (pink) around 14 min during steady-state inactivated hBEST1 current. *(B)* Repeated 1 min applications of 30 mM GABA during inactivation. In (A) and (B), the mean current at -100 mV (bottom) or +100 mV (top) from 250 ms voltage steps before and after the voltage ramp is shown. Zero current level is denoted by the dotted black line.

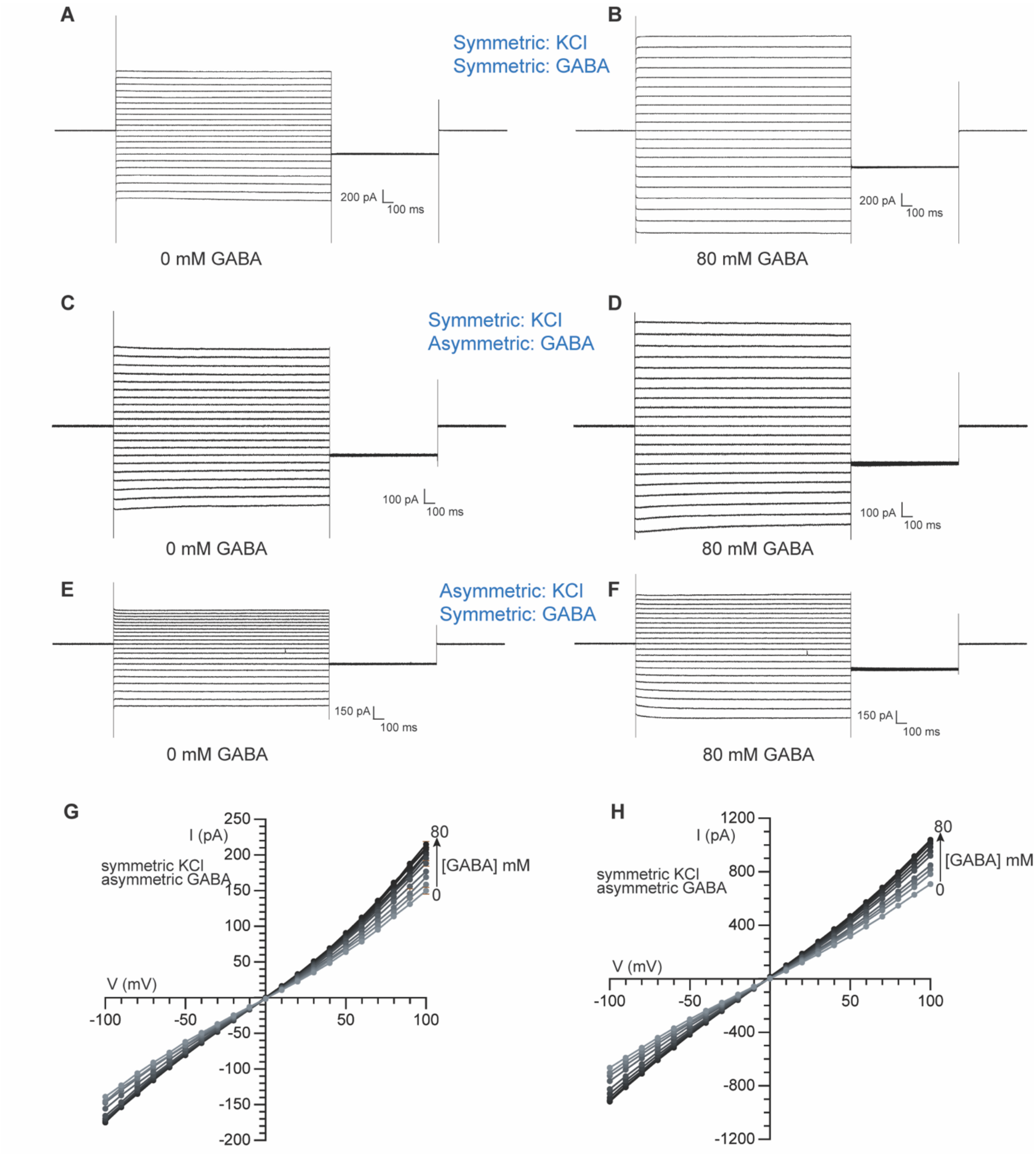


**Fig S3**. Planar lipid bilayer electrophysiology recordings of chicken BEST1. (*A*-*F*) Raw traces for the I-V relationships shown in Fig 3. The following protocol was used: holding potential of 0 mV for 0.5 s, then the voltage was stepped to values between −100 and +100 mV (with 10 mV increments) for 2 s. This was followed by a 1 s step to −40 mV. Finally, the voltage was returned to 0 mV. (*A*-*B*) Raw traces of 0 mM and 80 mM GABA used to calculate the I-V relationship at 0 and 80 mM shown in Fig 3*A*. (*C*-*D*) Raw traces of 0 mM and 80 mM GABA used to calculate the I-V relationship at 0 and 80 mM shown in Fig 3*C.* (*E*-*F*) Raw traces of 0 mM and 80 mM GABA used to calculate the I-V relationship at 0 and 80 mM shown in Fig 3*D*. (*G*-*H*) I-V relationships for experiments identical to that shown in Fig 3*C* (from different membranes).

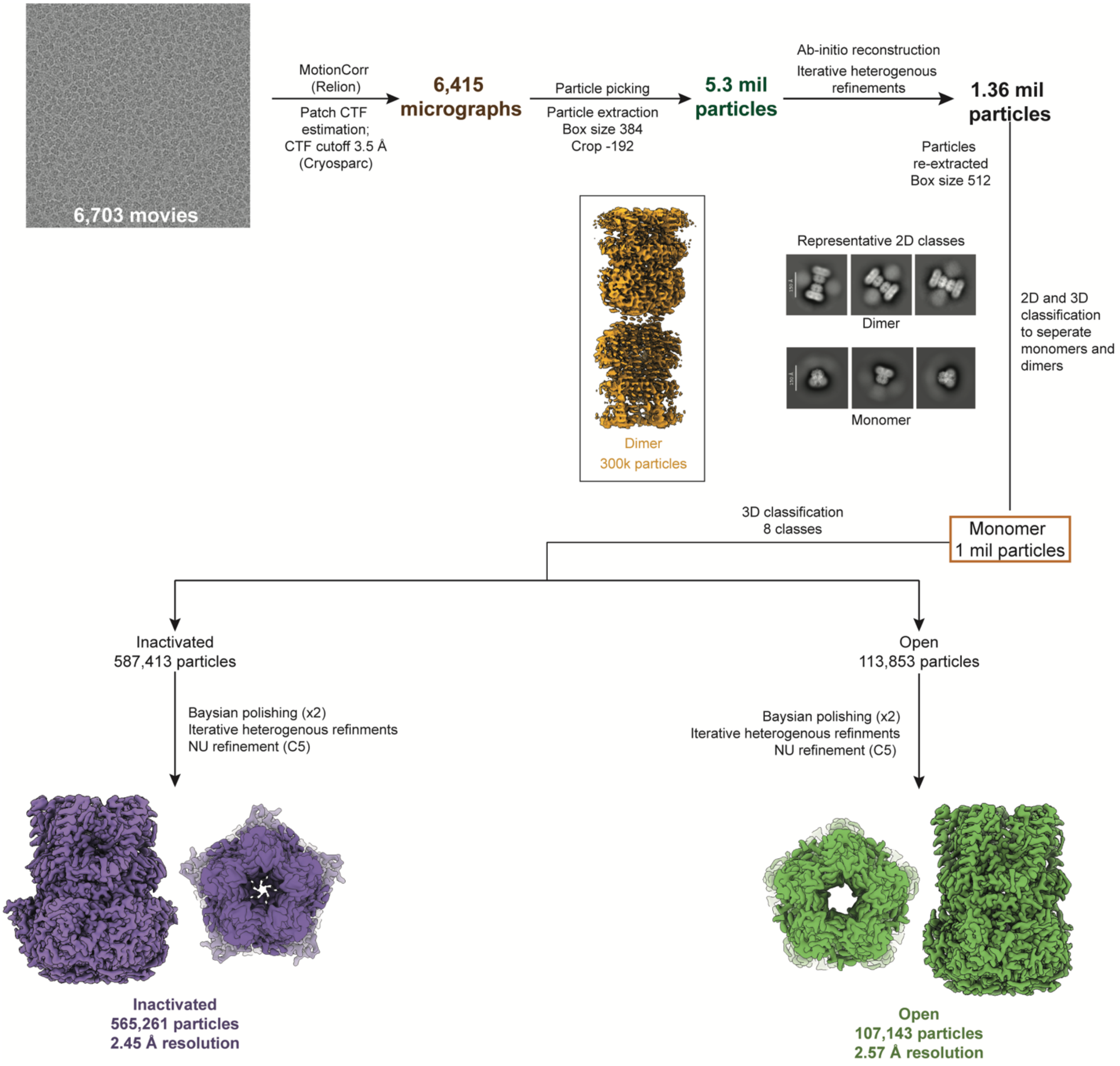


**Fig S4**. Cryo-EM data processing workflow for apo human BEST1 (without GABA). Details can be found in *Materials and Methods*.

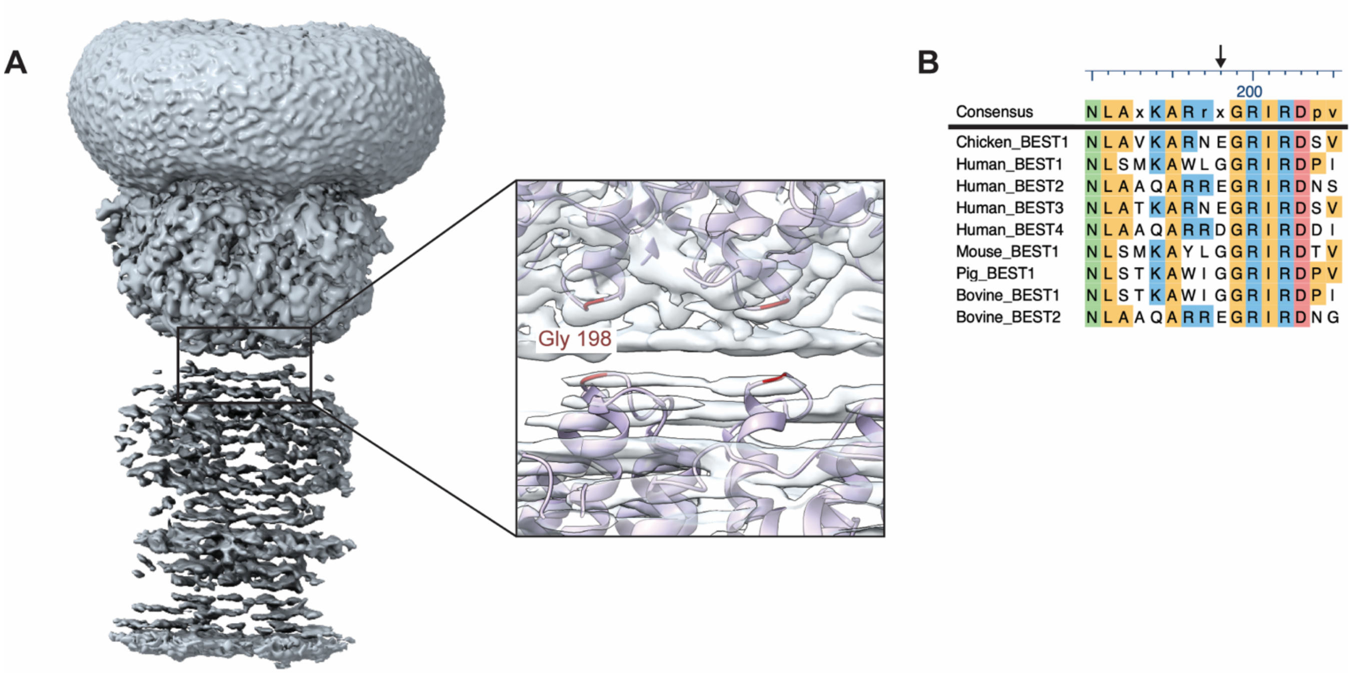


**Fig S5**. Dimer interface in human BEST1. *(A)* Dimer interface in the apo human BEST1 cryo-EM map is shown in the boxed region. Zoomed in view of the region shows that Gly198 (depicted in red) forms the dimer interface. *(B)* Sequence alignment of BEST1 paralogs. The position of amino acid 198 is denoted by an arrow.

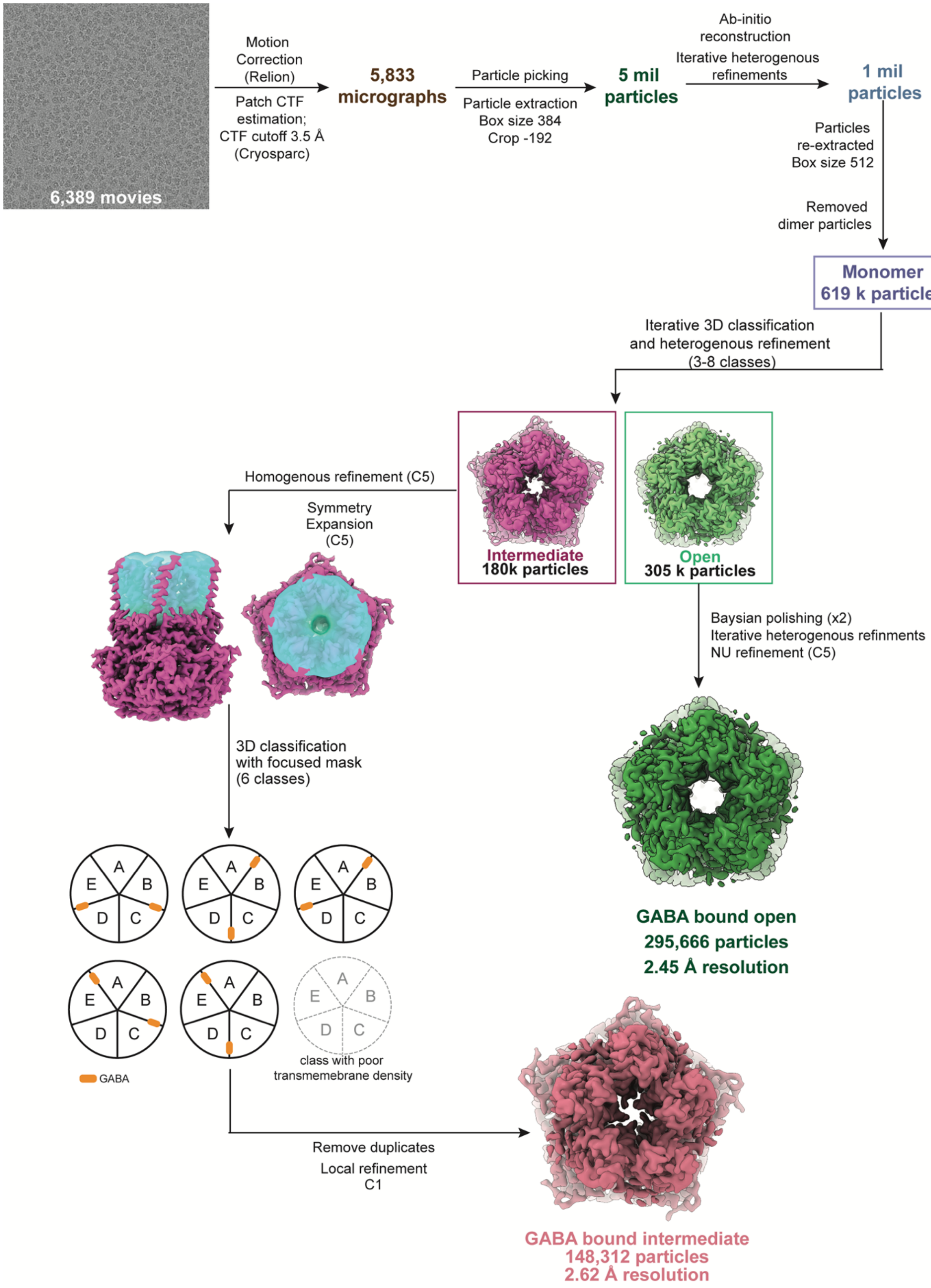


Fig S6. Cryo-EM data processing workflow for human BEST1 in complex with GABA. Details can be found in *Materials and Methods.*

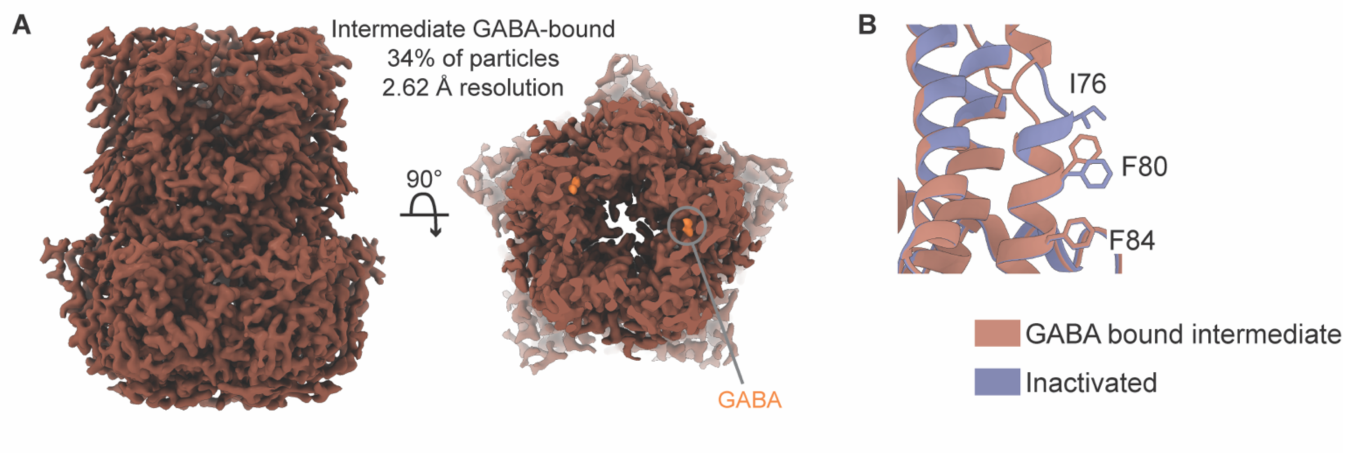


**Fig S7**. Structure of human BEST1 in an intermediate conformation with GABA bound. *(A)* 2.62 Å resolution cryo-EM map. Densities corresponding to GABA are colored orange (right panel). *(B)* GABA-bound intermediate structure compared to the inactivated structure (human BEST1). The neck region and GABA switch of the GABA-bound subunit is shown.

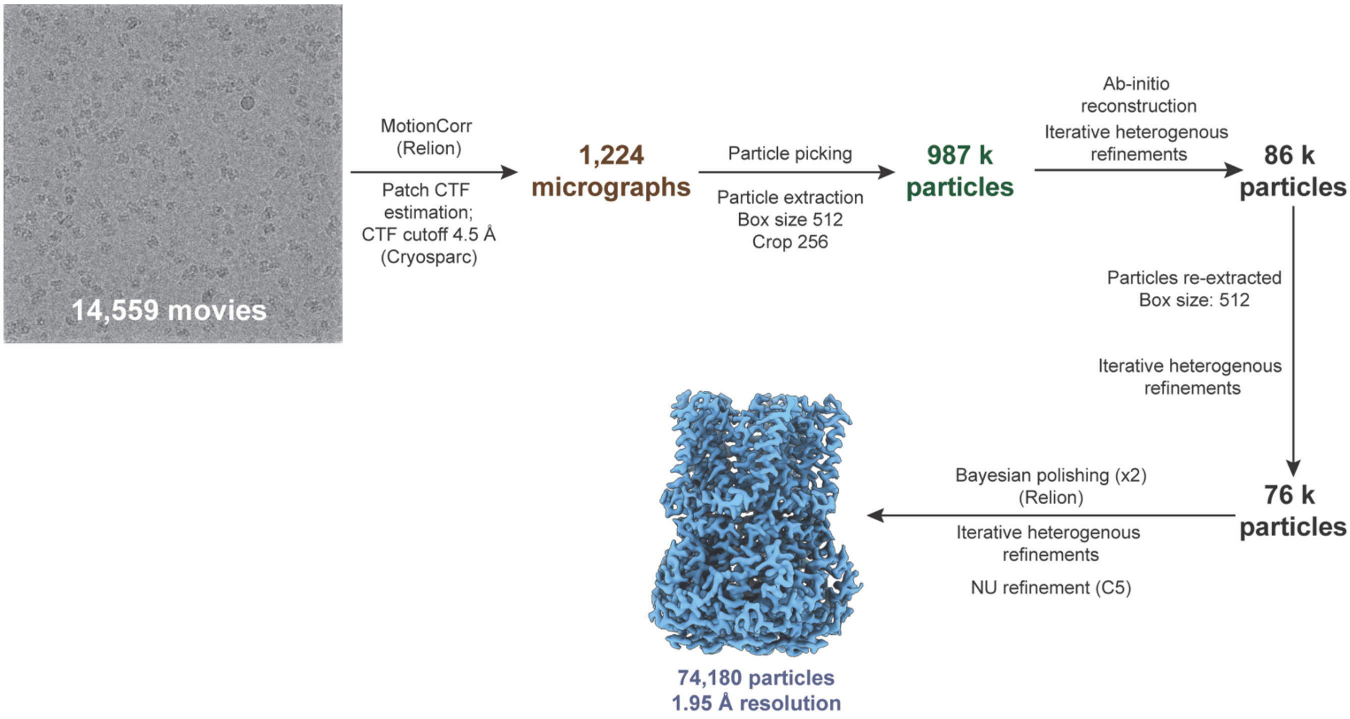


Fig S8. Cryo-EM data processing workflow for chicken BEST1 (1-345) in complex with GABA. Details can be found in *Materials and Methods*.

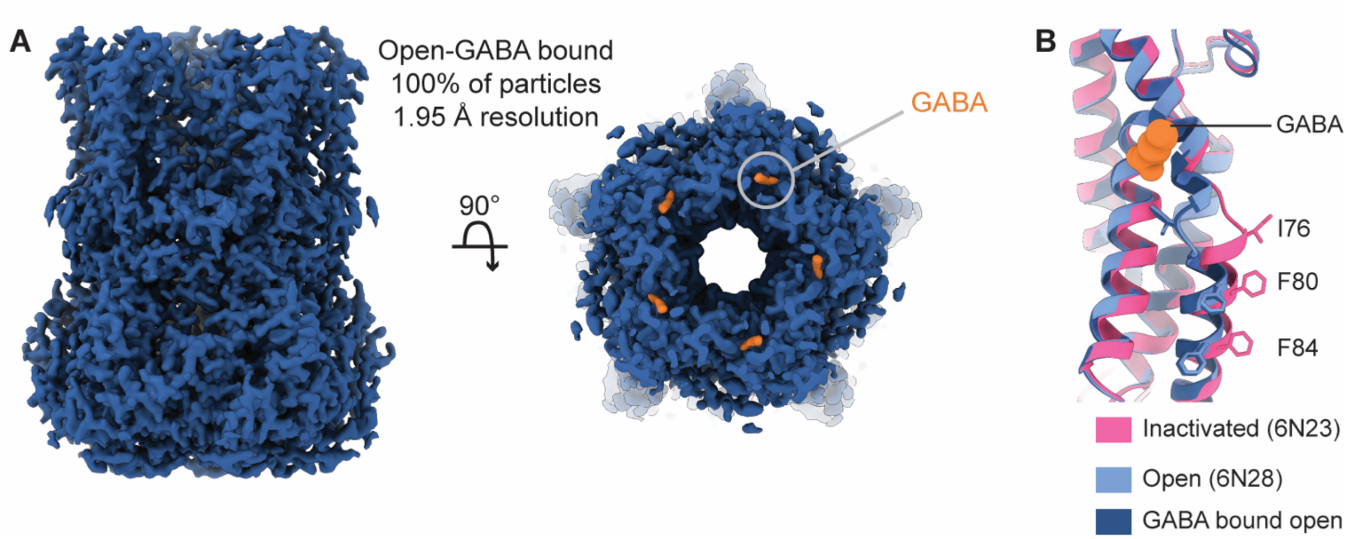


**Fig S9.** Cryo-EM structure of chicken BEST1 (1-345) in complex with GABA. *(A)* A 1.95 Å resolution map. Left, view from the side. Right, slice of the map from the extracellular side, showing a widened pore and densities corresponding to GABA (orange). *(B)* Comparison of a single subunit of the GABA-bound open structure (dark blue) to the previously determined structures of chicken BEST1 without GABA in an inactivated conformation (6N23; pink) and open conformation (6N28; blue).

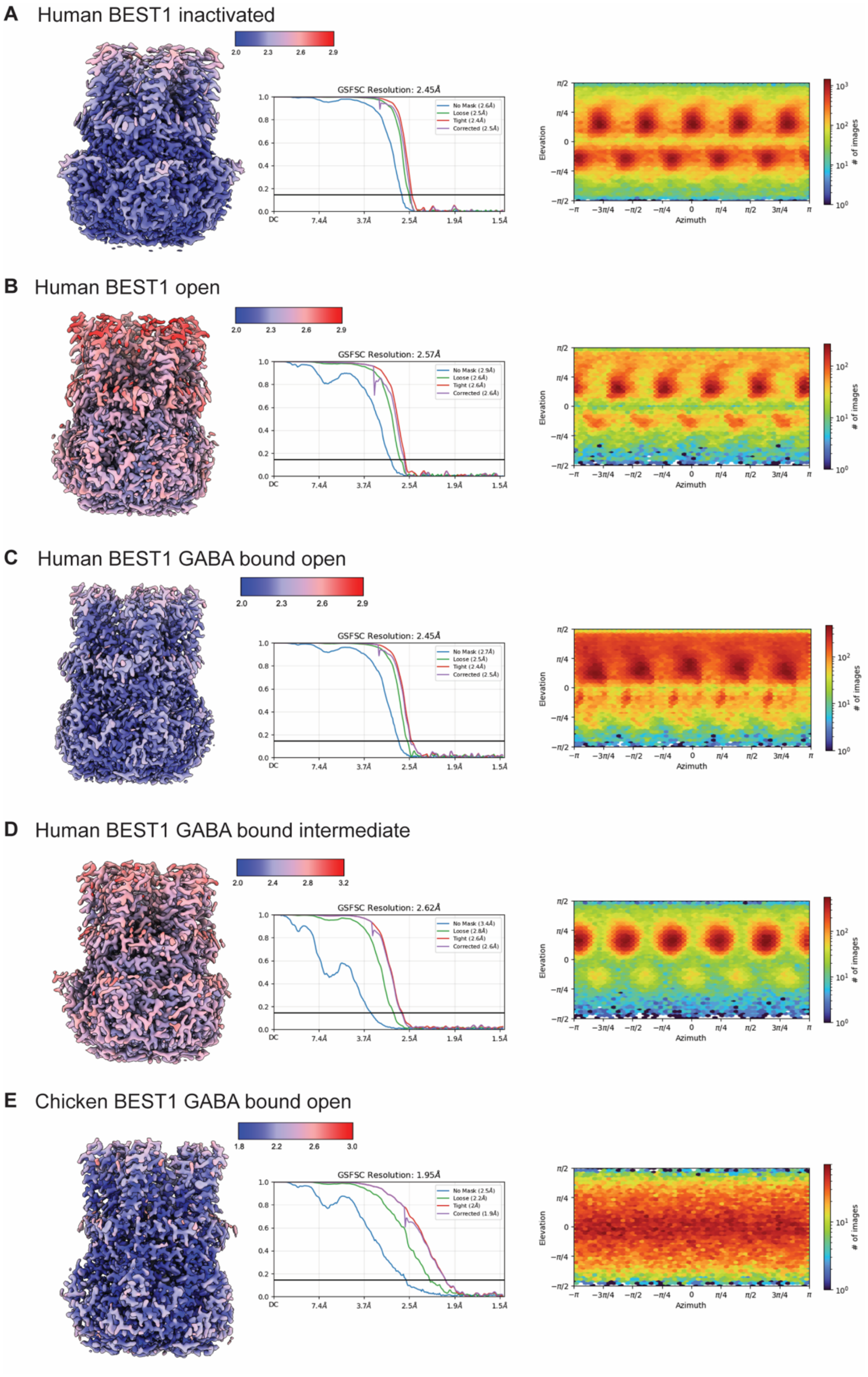


**Fig S10.** Cryo-EM map analyses, showing estimations of local resolutions, FSC curves, and the angular distributions of particles used in the final maps.

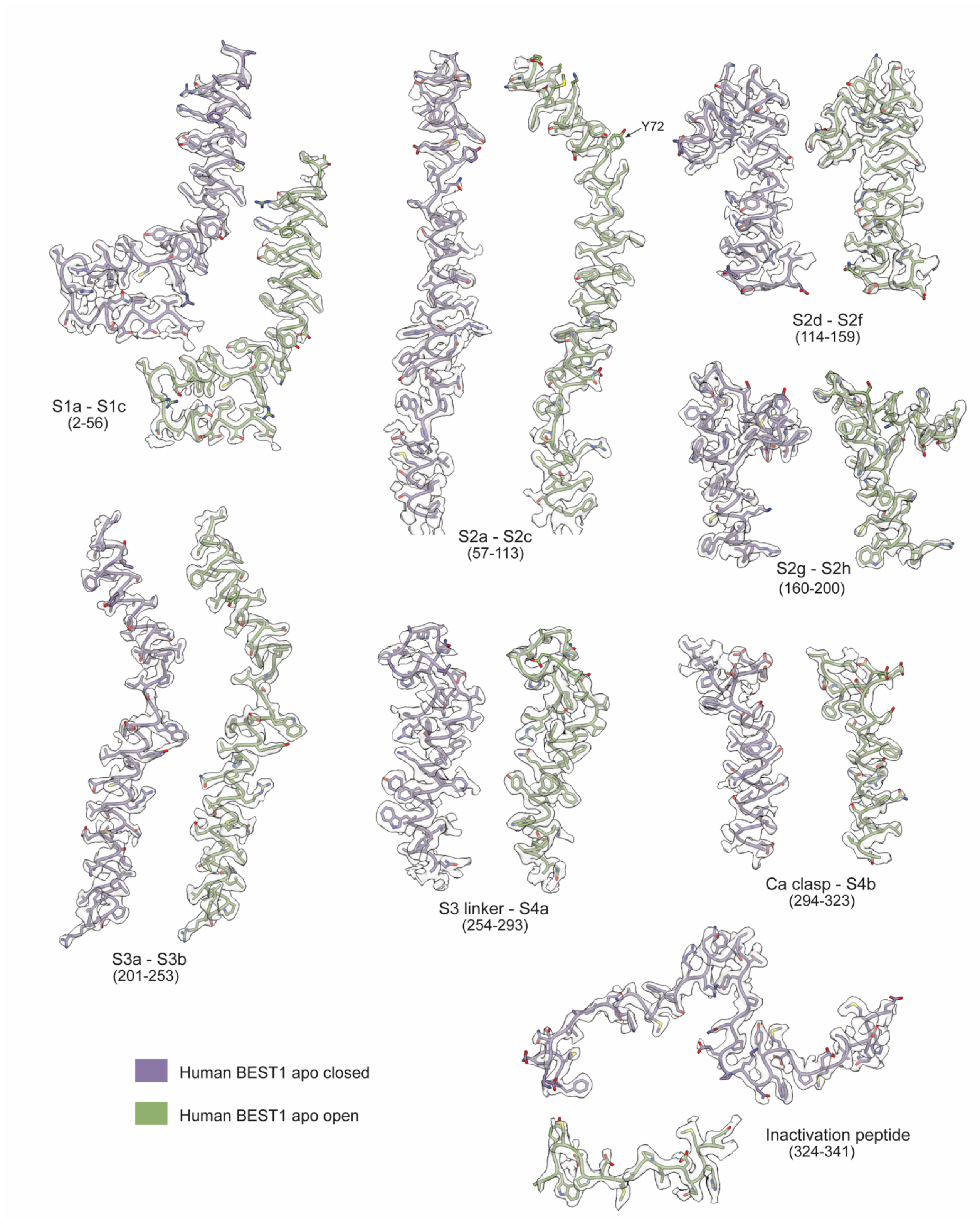


**Fig S11**. Cryo-EM densities of the human BEST1 structures without GABA that represent open and inactivated conformations. Densities (semi-transparent surface) for indicated regions are shown in the context of the atomic model (sticks).

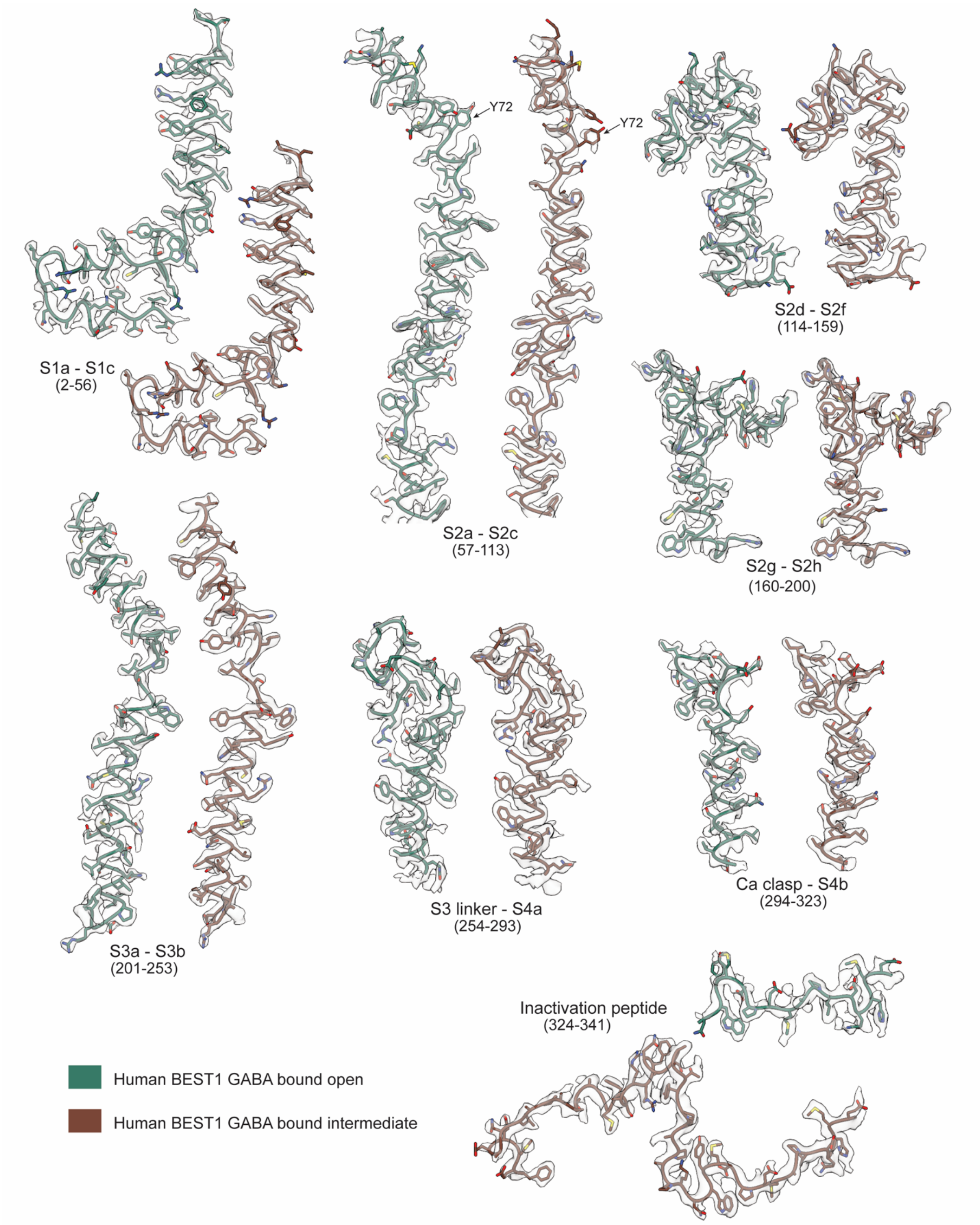


**Fig S12**. Cryo-EM densities of the human BEST1 structures with GABA that represent open and intermediate conformations (subunit without GABA bound). Densities (semi-transparent surface) for indicated regions are shown in the context of the atomic model (sticks).

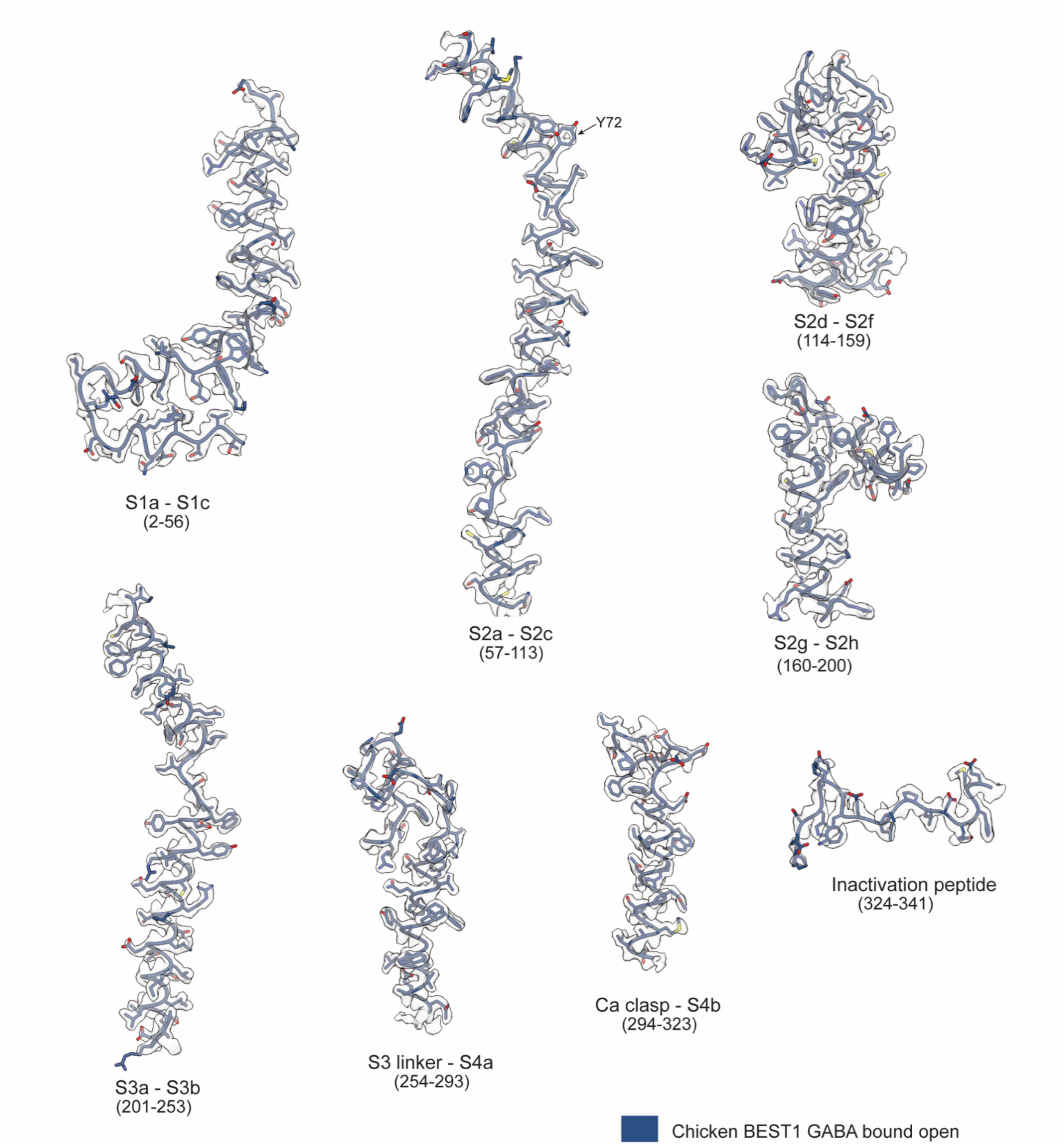


**Fig S13**. Cryo-EM density for chicken BEST1 (1-345) with GABA that represents an open conformation. Densities (semi-transparent surface) for indicated regions are shown in the context of the atomic model (sticks).

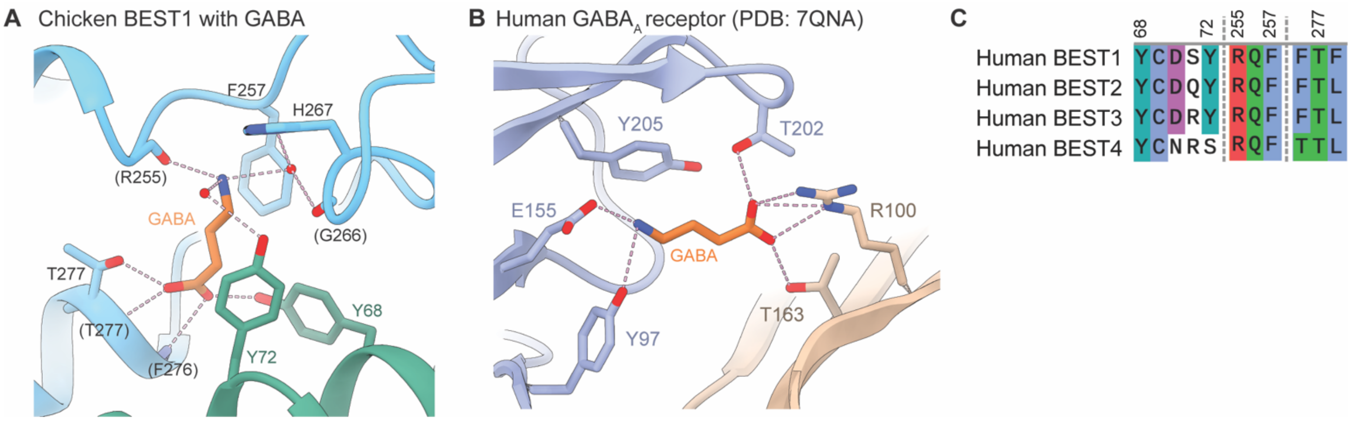


**Fig S14**. Comparison of the GABA binding site in BEST1 (chicken) with that of the GABA$_A$ receptor and sequence alignment. *(A)* The GABA binding site for chicken BEST1 is analogous to the site in human BEST1 (shown in **Fig** 6*D*). (B) The GABA binding site in a GABA$_A$ receptor (PDB: 7QNA). The $\beta$ subunit is shown in purple and the $\alpha$ subunit is colored beige. (C) Sequence alignment among human BEST1-4 for residues that coordinate GABA in BEST1 (human BEST1 amino acid numbering).

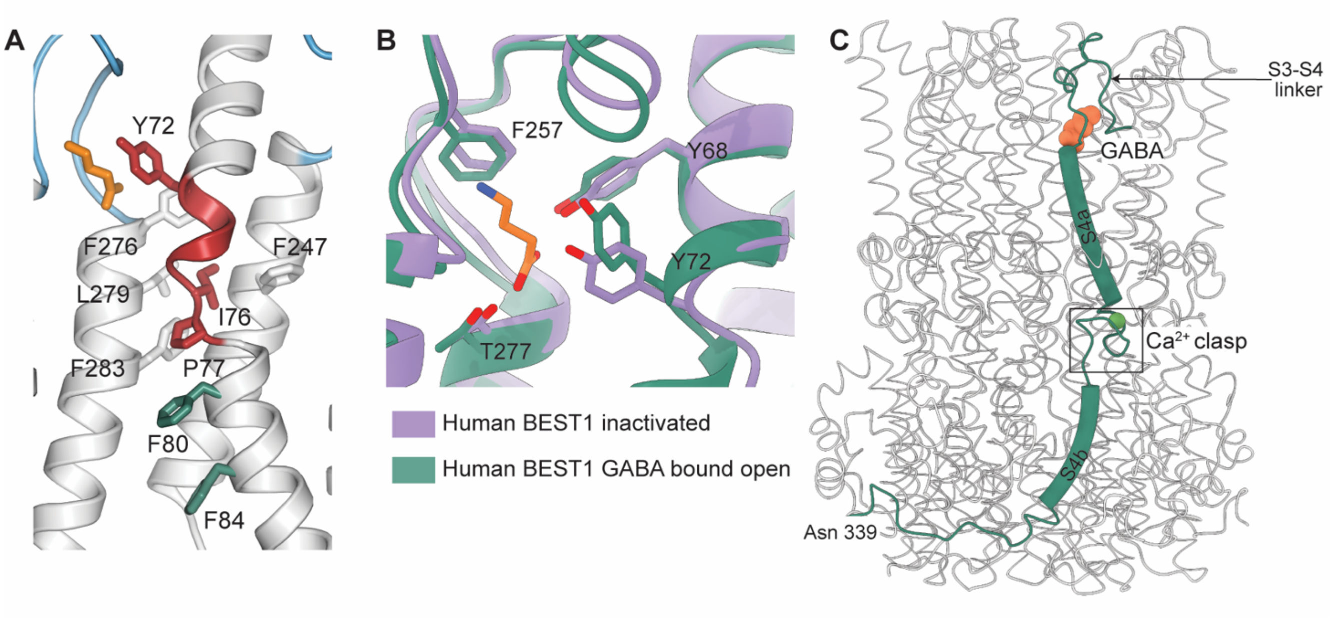


**Fig S15**. Binding pocket for Ile76 and location of the GABA site. *(A)* The binding pocket for Ile76 in the open conformation of the channel is shown, with amino acids that form van der Waals contacts drawn as sticks. GABA is shown as orange sticks. (*B*) Comparison of the conformations of amino acids comprising the GABA binding site from the inactivated structure (purple) and the GABA-bound open structure (green). GABA (from the GABA-bound open structure) is shown in orange. *(C)* GABA and $Ca^{2+}$ bind on opposite ends of the S4a helix. Select regions of one subunit are shown (green) with helices depicted as cylinders, in the context of a ribbon representation of the remainder of the channel (open conformation with GABA bound). A $Ca^{2+}$ ion is drawn as a green sphere. GABA is drawn as a sphere representation.

**Table S1**. Buffer composition for experiments in Fig 1.

| Figure panel | | Buffer |
|---|---|---|
| B | NaCl | 10 mM HEPES, pH 7.0, 0.2 mM EGTA, 0.1 mM $CaCl_2$, 2 μM ACMA, 0.5 mg/mL BSA, **65 mM NaCl and 50 mM $Na_2SO_4$** |
| | GABA | 10 mM HEPES, pH 7.0, 0.2 mM EGTA, 0.1 mM $CaCl_2$, 2 μM ACMA, 0.5 mg/mL BSA, **65 mM GABA and 50 mM $Na_2SO_4$** |
| | GABA + NaCl | 10 mM HEPES, pH 7.0, 0.2 mM EGTA, 0.1 mM $CaCl_2$, 2 μM ACMA, 0.5 mg/mL BSA, **65 mM NaCl and 65 mM GABA** |
| C | 0 | 10 mM HEPES, pH 7.0, 0.2 mM EGTA, 0.1 mM $CaCl_2$, 2 μM ACMA, 0.5 mg/mL BSA, **125 mM NaCl** |
| | 2 | 10 mM HEPES, pH 7.0, 0.2 mM EGTA, 0.1 mM $CaCl_2$, 2 μM ACMA, 0.5 mg/mL BSA, **125 mM NaCl and 2 mM GABA** |
| | 10 | 10 mM HEPES, pH 7.0, 0.2 mM EGTA, 0.1 mM $CaCl_2$, 2 μM ACMA, 0.5 mg/mL BSA, **125 mM NaCl and 10 mM GABA** |
| | 30 | 10 mM HEPES, pH 7.0, 0.2 mM EGTA, 0.1 mM $CaCl_2$, 2 μM ACMA, 0.5 mg/mL BSA, **125 mM NaCl and 30 mM GABA** |
| D | Control | 10 mM HEPES, pH 7.0, 0.2 mM EGTA, 0.1 mM $CaCl_2$, 2 μM ACMA, 0.5 mg/mL BSA, **125 mM NaCl** |
| | Glycine | 10 mM HEPES, pH 7.0, 0.2 mM EGTA, 0.1 mM $CaCl_2$, 2 μM ACMA, 0.5 mg/mL BSA, **125 mM NaCl and 30 mM Glycine** |
| | D-serine | 10 mM HEPES, pH 7.0, 0.2 mM EGTA, 0.1 mM $CaCl_2$, 2 μM ACMA, 0.5 mg/mL BSA, **125 mM NaCl and 30 mM D-serine** |
| | β-Alanine | 10 mM HEPES, pH 7.0, 0.2 mM EGTA, 0.1 mM $CaCl_2$, 2 μM ACMA, 0.5 mg/mL BSA, **125 mM NaCl and 30 mM β-Alanine** |
| | GABA | 10 mM HEPES, pH 7.0, 0.2 mM EGTA, 0.1 mM $CaCl_2$, 2 μM ACMA, 0.5 mg/mL BSA, **125 mM NaCl and 30 mM GABA** |
| E | Control | 10 mM HEPES, pH 7.0, 0.2 mM EGTA, 0.1 mM $CaCl_2$, 2 μM ACMA, 0.5 mg/mL BSA, **125 mM NaCl and 15 mM $Na_2SO_4$** |
| | Gluconate | 10 mM HEPES, pH 7.0, 0.2 mM EGTA, 0.1 mM $CaCl_2$, 2 μM ACMA, 0.5 mg/mL BSA, **125 mM NaCl and 30 mM NaGluconate** |
| | Glutamate | 10 mM HEPES, pH 7.0, 0.2 mM EGTA, 0.1 mM $CaCl_2$, 2 μM ACMA, 0.5 mg/mL BSA, **125 mM NaCl and 30 mM NaGlutamate** |
| | GABA | 10 mM HEPES, pH 7.0, 0.2 mM EGTA, 0.1 mM $CaCl_2$, 2 μM ACMA, 0.5 mg/mL BSA, **125 mM NaCl, 15 mM $Na_2SO_4$, 30 mM GABA** |
| F | NaCl | 5 mM HEPES, pH 7.0, 200 μM $CaCl_2$, 2 μM ACMA, 0.5 mg/mL BSA, **65 mM NaCl and 50 mM $Na_2SO_4$** |
| | GABA | 5 mM HEPES, pH 7.0, 200 μM $CaCl_2$, 2 μM ACMA, 0.5 mg/mL BSA, **65 mM GABA and 50 mM $Na_2SO_4$** |
| | GABA + NaCl | 5 mM HEPES, pH 7.0, 200 μM $CaCl_2$, 2 μM ACMA, 0.5 mg/mL BSA, **65 mM NaCl and 65 mM GABA** |

**Table S2**. Data collection, refinement, and validation statistics.

| | Human BEST1 inactivated PDB: 9EGS EMDB: EMD-47996 | Human BEST1 open PDB: 9EGT EMDB: EMD-47997 | Human BEST1 GABA bound open PDB: 9EGM EMDB: EMD-47991 | Human BEST1 GABA bound intermediate PDB: 9EGQ EMDB: EMD-47995 | Chicken BEST1 GABA bound open PDB: 9EFZ EMDB: EMD-47982 |
|---|---|---|---|---|---|
| Data collection and processing | | | | | |
| Microscope | FEI Titan Krios (MSKCC) | FEI Titan Krios (MSKCC) | FEI Titan Krios (MSKCC) | FEI Titan Krios (MSKCC) | FEI Titan Krios (NYSBC) |
| Camera | Falcon 4i | Falcon 4i | Falcon 4i | Falcon 4i | Falcon 4i |
| Magnification | 165,000× | 165,000× | 165,000× | 165,000× | 165,000× |
| Voltage (kV) | 300 | 300 | 300 | 300 | 300 |
| Electron exposure ($e^{-}/Å^2$) | 60.30 | 60.30 | 60.08 | 60.08 | 43.70 |
| Defocus range (μm) | −1.0 ~ −2.0 | −1.0 ~ −2.0 | −1.0 ~ −2.0 | −1.0 ~ −2.0 | -0.6 ~ -2 |
| Pixel size (Å) | 0.725 | 0.725 | 0.725 | 0.725 | 0.725 |
| Software | RELION 3.1 cryoSPARC v4.5.0 | RELION 3.1 cryoSPARC v4.5.0 | RELION 3.1 cryoSPARC v4.5.0 | RELION 3.1 cryoSPARC v4.5.0 | RELION 3.1 cryoSPARC v4.5.0 |
| Symmetry imposed | C5 | C5 | C5 | C1 | C5 |
| Initial particle images (no.) | 5,338,248 | 5,338,248 | 4,996,553 | 4,996,553 | 987,499 |
| Final particle images (no.) | 565,261 | 107,143 | 295,666 | 148,312 | 74,180 |
| Overall map resolution (Å) FSC threshold 0.143 | 2.45 | 2.57 | 2.45 | 2.62 | 1.95 |
| Refinement | | | | | |
| Software | Phenix 1.21 real-space-refine | Phenix 1.21 real-space-refine | Phenix 1.21 real-space-refine | Phenix 1.21 real-space-refine | Phenix 1.21 real-space-refine |
| Initial model (PDB code) | | | | | 6N28 |
| Model resolution (Å) FSC threshold 0.5 | 2.5 | 2.6 | 2.5 | 2.7 | 2.0 |
| Map sharpening B factor ($Å^2$) | -65 | -65 | -80 | -84 | -38 |
| Model composition | | | | | |
| Non-hydrogen atoms | 15805 | 14169 | 14255 | 15267 | 14477 |
| Protein residues | 1885 | 1690 | 1690 | 1875 | 1700 |
| Ligands | $Ca^{2+}$: 5 $Cl^{-}$: 10 | $Ca^{2+}$:5 | $Ca^{2+}$: 5 $Cl^{-}$: 5 GABA:5 | $Ca^{2+}$: 5 GABA: 2 | $Ca^{2+}$: 5 $Cl^{-}$: 5 GABA: 5 |
| Water | 270 | 269 | 330 | 0 | 382 |
| Mean B factors ($Å^2$) | | | | | |
| Protein | 12.17 | 20.59 | 15.58 | 38.21 | 24.76 |
| Ligands/Water | 22.91/15.11 | 16.45/20.32 | 23.64/12.45 | 41.68 | 42.70/26.56 |
| R.m.s deviations | | | | | |
| Bond lengths (Å) | 0.003 | 0.003 | 0.003 | 0.005 | 0.002 |
| Bond angles (°) | 0.516 | 0.507 | 0.494 | 0.647 | 0.419 |
| Validation | | | | | |
| MolProbity score | 1.58 | 1.47 | 1.45 | 1.43 | 1.26 |
| Clashscore | 10.73 | 8.76 | 8.18 | 7.89 | 4.89 |
| Poor rotamers (%) | 0.9 | 0.34 | 1.01 | 0.62 | 0.32 |
| Ramachandran plot | | | | | |
| Favored (%) | 97.87 | 98.81 | 98.51 | 98.82 | 99.41 |
| Allowed (%) | 2.13 | 1.19 | 1.49 | 1.18 | 0.59 |
| Disallowed (%) | 0 | 0 | 0 | 0 | 0 |
